# Community diversity determines the evolution of synthetic bacterial communities under artificial selection

**DOI:** 10.1101/2021.09.14.460260

**Authors:** Tiffany Raynaud, Marion Devers, Aymé Spor, Manuel Blouin

**Affiliations:** Agroécologie, AgroSup Dijon, INRAE, Univ. Bourgogne, Univ. Bourgogne Franche-Comté, F-21000 Dijon, France

**Keywords:** artificial selection efficiency, community diversity, community evolution, synthetic bacterial communities

## Abstract

Artificial selection can be conducted at the community level in the laboratory through a differential propagation of the communities according to their level of expression of a targeted function (i.e. community phenotype). Working with communities instead of individuals as selection units brings in additional sources of variation in the considered phenotype that can arise through changes in community structure and influence the outcome of the artificial selection. These sources of variation could even be increased by manipulating species diversity. In this study, we wanted to assess the effect of manipulating initial community richness on artificial selection efficiency, defined as the change in the targeted function over time as compared to a control treatment without artificial selection. We applied artificial selection for a high productivity on synthetic bacterial communities varying for their initial richness level (from one to 16 strains). Our results showed that, overall, the communities that were artificially selected were 16% more productive than the control communities. Community richness positively influenced community productivity and metabolic capacities and was a strong determinant of the dynamics of community evolution. Our results suggested that community richness could influence artificial selection efficiency but a convergence of the community composition might have limited the effect of diversity on artificial selection efficiency. We propose that applying artificial selection on communities varying for their diversity could allow to find communities differing for their level of expression of a function but also for their responsiveness to artificial selection, provided that their initial composition is different enough.

## Introduction

In 1989, in the framework of the levels-of-selection theory (or multi-level selection), Wilson and Sober supported the idea that natural selection at the community level could occur *in natura*. In the two last decades, several attempts have been made to enhance or reduce a property or a function performed by a microbial community through artificial selection at the community level in the laboratory. The general principle is to *i*) grow replicates of a microbial community, *ii*) assess the performance of these communities regarding the targeted function, *iii*) propagate the best performing community(ies). This approach has been used to study the degradation of a toxic compound (Swenson et al. 2000a), the modification of the pH of an aquatic medium (Swenson et al. 2000b), CO_2_ emissions (Blouin et al. 2015), chitinase activity (Wright et al. 2019), productivity (Raynaud et al. 2019), the hydrolysis of starch (Chang et al. 2020) and the growth promotion of a bacterial strain (Chang et al. 2020). All of the studies involving artificial selection of microbial communities showed that the engineering of complex microbial communities is not straightforward. In particular, improvements of the targeted function are often unstable (Swenson et al. 2000b; Raynaud et al. 2019; Wright et al. 2019) and registered at the beginning of the procedure only (Chang et al. 2020). This might be related to a decrease in community phenotypic variance and heritability over time (Blouin et al. 2015) but also to changes in community structure, due to the succession of species for example (Wright et al. 2019), that could limit artificial selection efficiency.

Artificial selection can be applied without any *a priori* knowledge of community composition or functioning but, assessing community diversity during an artificial selection experiment can allow a better understanding of community evolutionary dynamics in this context. It is well-known that the components of the diversity of a community (e.g. community richness, composition, evenness) can have an influence on many functions such as productivity or stability (Hooper et al. 2005). Several studies, conducted on bacterial communities and manipulating community richness (up to 72 species in Bell et al. (2005)), experimentally tested for effects of community diversity on the community respiration rate (Bell et al. 2005) or productivity (Gravel et al. 2011; Fetzer et al. 2015). These studies highlighted a positive and saturating relationship between the increase in community richness and the increase in the level of the measured function. Two main categories of mechanism can underlie a diversity-function relationship (Loreau et al. 2001): complementarity effects (i.e. the function is due to a combination of species through niche partitioning or facilitation between species) and selection effects (i.e. the function is due to a dominant species). Increasing community richness increases the probability of these mechanisms to occur (Loreau and Hector 2001) and thus to observe an increase in the studied function.

Beyond the effect of community diversity on the initial level of a function, increasing community diversity could also influence community response to selection. The term “evolution” is sometimes restricted to genetic changes over generations (Barraclough 2015). But, when it comes to community evolution, additional sources of variations can be involved in the community evolutionary response (Penn 2003; Williams and Lenton 2007), provided that they can be transmitted to the next “generation of communities” (i.e. that they are heritable; Goodnight 2000). Indeed, the community phenotype can result from allelic composition and intragenomic interactions (i.e. epistasis), population composition and intraspecific interactions, and from species composition and interspecific interactions. All these sources of variations in community response to artificial selection depend on community diversity. Selecting at the community level while increasing species richness and thus the different sources of variations may increase the probability to observe extreme values for the targeted function among the fixed number of communities under selection. This increasing number of species should thus increase the selection differential (S) in the breeder equation (R = h^2^ x S, with R the response to selection and h^2^ the heritability; Lush 1937). As a consequence, the response to selection (R) should be higher when there are many, as compared with few species, provided that the phenotype is reliably transmitted between parent and offspring communities (i.e. h^2^ > 0).

In this study, we wanted to explore the link between the diversity of a community and the efficiency of artificial selection. Previous artificial selection experiments were mainly conducted on complex natural microbial communities (retrieved from soil or plant leaves for example) that were then grown under laboratory conditions. Some studies assessed the changes in microbial community diversity over the course of the experiment (Raynaud et al. 2019; Jacquiod et al. 2021) but community diversity was not intentionally manipulated. We designed an experiment that combined the approaches developed in artificial selection of microbial communities and in biodiversity-ecosystem functioning experiments. An artificial selection procedure was applied on synthetic bacterial communities including five richness levels from one to 16 strains. We defined artificial selection efficiency as the change in the targeted phenotype – a high productivity – over time as compared to a control treatment without artificial selection. We hypothesized that increasing the diversity of the selected communities could be responsible for a larger range of variation in productivity, providing more opportunities for selection to act, and thus enhancing the efficiency of artificial selection.

## Materials and methods

### Bacterial strains

Eighteen bacterial strains were used in this experiment. They were chosen based on the screening of 38 laboratory strains for their ability to grow in the chosen experimental conditions (detailed below). Based on the growth curves of the 38 strains (assessed by Bioscreen, Oy Growth Curves Ab Ltd, Finland), we excluded the strains that showed an absence of growth, a slow growth or a decline, as well as strains that had a longer lag phase or a faster growth than the others. This was done to avoid the dominance of one or few strains from the very beginning of the experiment in communities due to too large differences in growth ability in our culture conditions. The 18 chosen strains belonged to three phyla, six classes and 12 genera (Table 1).

**Table 1:** Bacterial strains used in the experiment. Note that *Aminobacter aminovorans* was previously known as *Chelatobacter heintzii*.

| Strain | Phylum | Class |
| --- | --- | --- |
| *Alcaligenes eutrophus* JMP131 | Proteobacteria | β-proteobacteria |
| *Agrobacterium* sp. 9023 | Proteobacteria | α-proteobacteria |
| *Aminobacter aminovorans* SR38 | Proteobacteria | α-proteobacteria |
| *Arthrobacter* sp. | Actinobacteria | Actinobacteria |
| *Arthrobacter* sp. BS2 | Actinobacteria | Actinobacteria |
| *Cupriavidus necator* JMP134 | Proteobacteria | β-proteobacteria |
| *Dyadobacter fermentans* DSM 18053 | Bacteroidetes | Flavobacteria |
| *Escherichia coli* K12 | Proteobacteria | γ-proteobacteria |
| *Escherichia coli* WA803 | Proteobacteria | γ-proteobacteria |
| *Microbacterium* sp. C448 | Actinobacteria | Actinobacteria |
| *Pseudomonas azelaica* | Proteobacteria | γ-proteobacteria |
| *Pseudomonas knackmussii* DSM 6978 | Proteobacteria | γ-proteobacteria |
| *Pseudomonas* sp. ADP3 | Proteobacteria | γ-proteobacteria |
| *Pseudomonas* sp. ADPe | Proteobacteria | γ-proteobacteria |
| *Pseudopedobacter saltans* DSM 12145 | Bacteroidetes | Sphingobacteria |
| *Ralstonia* sp. | Proteobacteria | β-proteobacteria |
| *Sphingomonas wittichii* RW1 | Proteobacteria | α-proteobacteria |
| *Variovorax* sp. 38R | Proteobacteria | β-proteobacteria |

### Community construction

The 18 strains were grown under five levels of initial richness: 1, 2, 4, 8, 16 strains. All the monocultures were grown (i.e. n=18 for level 1), and six communities per remaining level of richness were established (i.e. n=6 for levels 2, 4, 8 and 16). Community composition was determined by randomly choosing 16 strains among 18 without replacement to construct the six communities of the richness level 16. We built the communities from the lower richness levels as subsets of the communities of the upper richness level. The first eight strains that were randomly assigned to the first community of richness level 16 composed the first community of level 8 and so on until creating six communities of eight strains. The same method was used for levels 4 and 2.

### Growth conditions

The culture medium was a mix of 1:5 lysogeny broth (LB) and 1:5 tryptic soy broth (TSB), these media notably differ for their carbon sources which can allow for niche partitioning (Van den Bergh et al. 2018). Before the start of the experiment, each strain was grown on a Petri dish (1:5 LB+TSB with agar) by streaking, starting from monocultures stored in 30% glycerol at -80 °C. Then, one colony per strain was picked and placed in 42 ml of culture medium in a flask (28 °C, 110 rpm, 24 h). The optical density (OD) was assessed at 600 nm (BioPhotometer 6131, Eppendorf, Germany) and each suspension was diluted to a final OD of 0.002. The diluted suspensions were used to inoculate monocultures and to build the communities by adding an equivalent volume of each required suspension according to the composition of the community. The monocultures and communities were grown in sterile 2 ml deep-well plates (Porvair Sciences, UK) filled with 1 ml of culture medium. Each monoculture (n=18) and community (n=24) was replicated 11 times i.e. 11 wells of a plate were inoculated with the same suspension. It resulted in six plates in total over which each level of richness was represented and an extra suspension was added as a control to track for possible “plate effects”. Temperature was kept to 28°C and there was no shaking in order to allow for possible spatial niche partitioning (Van den Bergh et al. 2018).

### Experimental evolution

A transfer into a new plate and fresh medium occurred every 84 h for 20 weeks resulting in 40 artificial selection cycles. The phenotype targeted by artificial selection was a high productivity which was assessed by OD measurement. Each 84 h, the content of the wells was homogenised by pipetting up and down and 200 μL of suspension were transferred into a microplate-reader compatible plate (Fisherbrand 96-Well plates, Fisher Scientific, USA). The OD was measured at 600 nm (Infinite M200 PRO, Tecan, Switzerland) and the 200 μL of suspension were discarded. The artificial selection treatment (AS) occurred through *i*) the identification of the well among ten showing the highest OD, *ii*) the sampling of 20 μL of the corresponding suspension and *iii*) the inoculation of 980 μL of fresh medium with these 20 μL of suspension. The two latter steps were repeated until the ten wells of the new plate were inoculated. As previously mentioned, there were 11 wells for each monoculture or community; while ten wells were dedicated to the artificial selection, the remaining well was used as a control without artificial selection. 20 μL of suspension were sampled from this well and inoculated into 980 μL of fresh medium whatever the OD of the suspension (No artificial Selection, NS). This treatment corresponded to a controlled natural selection (Conner 2003) in which environmental conditions were imposed but the communities were allowed to reproduce without artificial selection. At each transfer event, the suspensions that were used to inoculate the new plate (i.e. the suspension from the selected wells in AS and the suspension from the control wells) were stored in 30% glycerol at -80°C.

### Post-selection

After the end of the experimental evolution, we revived the monocultures and communities from cycle 0 (i.e. initial inocula, hereafter called “ancestors”) and 40 (i.e. cultures after 40 selection cycles, hereafter called “evolved under AS”) from glycerol stocks. The control cultures (evolved under NS) were also included. The aim was to assess the phenotype of ancestors and evolved monocultures and communities at the same time in one experiment to corroborate what was observed in the artificial selection experiment. The monocultures and communities were first grown in 20 ml of culture medium in flasks (28°C, 110 rpm, 24 h for the evolved under AS and NS and 48 h for the ancestors as 24 h were not enough for them to reach sufficient OD). The optical density was assessed at 600 nm (Infinite M200 PRO, Tecan, Switzerland), each suspension was diluted in culture medium to a final OD of equivalent 0.002 in BioPhotometer 6131 and allowed to grow in triplicate for 84 h in the growth conditions of the experimental evolution (i.e. deep-well plates, 28°C, no shaking, 1 ml of 1:5 LB+TSB). The OD was measured at 600 nm after 84 h of growth as previously described (see “Experimental evolution”). Three bacterial strains were grown on each plate (six in total) as controls for a possible “plate effect”.

### Description of the growth dynamics

To go further in the phenotypic description of the ancestor and evolved monocultures and communities, we used the same protocol as previously described (culture in flask at 110 rpm, 28 °C, for either 24 or 48 h, OD measurement, dilution to a final OD of 0.002) and inoculated triplicate of the suspensions into microplates that were suitable for the detailed analysis of growth kinetics (sterile plates honeycomb, Thermo Scientific, USA). The growth conditions were: 200 μL of 1:5 LB+TSB, 28 °C, 15 s of shaking occurring 5 s before each OD measurement, one measurement every 30 min for 84 h (Bioscreen, Oy Growth Curves Ab Ltd, Finland). Three bacterial cultures were grown on each plate (seven in total) as controls for a possible “plate effect”.

### Metabolism

We assessed the metabolic capabilities of the ancestor and evolved monocultures and communities (under AS and NS) using EcoPlates (Biolog, USA). 31 carbon (and nitrogen) sources belonging to six categories (amino acids, amines, carbohydrates, carboxylic acids, phenolic compounds and polymers ; Montserrat Sala et al. 2010) were tested. In the same way as the post-selection experiment and the growth dynamic description experiment, ancestors and evolved (under AS and NS) monocultures and communities were revived and grown in flasks (110 rpm, 28°C, either 24h or 48h). Then, OD was assessed and the suspensions were diluted in 0.9% NaCl solution to reach a final OD of 0.2 (tests were conducted before the start of the experiment and showed that cell washing gave similar results to those obtained with a dilution approach indicating that a remaining amount of culture medium in the suspension did not changed the results). EcoPlates were inoculated with 120 μL of diluted suspension (one plate per sample, each substrate was repeated three times per plate) and placed at 28 °C without shaking. When a substrate was used by the bacteria, a tetrazolium dye was reduced which produced a purple coloration which was assessed by OD measurement at 590 nm (Infinite M200 PRO, Tecan, Switzerland). A first measurement was done four hours after the inoculation and then twice a day for four days.

### Community composition analysis

To assess for changes in community composition over the experiment, we performed 16S rRNA gene and *gyrB* sequencing on communities from selection cycles 1, 14, 27 and 40 for both AS and NS. In order to track for the presence of contaminants, 16S rRNA gene and *gyrB* sequencing was performed on monocultures from selection cycles 1 and 40 (see Appendix 1 for the details on DNA extractions, PCR and bioinformatics analyses).

### Statistical analyses

We first ran preliminary analyses to check for the presence of contaminants in the samples (i.e. strains that were not included in the initial species composition) and to determine at which step of the experimentations they occurred as it could have or not an influence on the results and more particularly on the validation of our main hypothesis (see Appendix 2). It resulted in the removal of 16.7% of the samples in the experimental evolution dataset and 14.3% of the samples in the growth dynamics, metabolism and post-selection datasets. The smallest sample size (n) occurred for the artificial selection treatment at richness levels 2 and 4 for which n was equal to 4 (instead of 6). The following analyses were conducted on the resulting datasets.

The data from the experimental evolution procedure were analysed with the following linear mixed model:

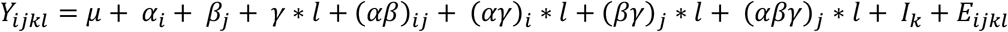

*Y*_*ijkl*_ is the OD of the individual (i.e. monoculture or community) of identity *k*, of initial richness level *i*, under selection treatment *j*, at cycle *l*. *µ* is the intercept. *α*_*i*_ is the effect of the initial richness level (qualitative: 1, 2, 4, 8, 16), *β*_*j*_ is the effect of the selection treatment (qualitative: AS, NS). *γ* ∗ *l* is the effect the selection cycle (quantitative: 1 to 40). The interaction effects between *i*) the initial richness level and the selection treatment (*αβ*)_*ij*_; *ii*) the initial richness level and the selection cycle (*αγ*)_*i*_ ∗ *l*; *iii*) the selection treatment and the selection cycle (*βγ*)_*j*_ ∗ *l*; *iv*) the initial richness level, the selection treatment and the selection cycle (*αβγ*)_*j*_ ∗ *l* were also included in the model. *I*_*k*_ is the random effect of the individual and *E*_*ijkl*_ is the residual error. An autoregression structure of order 1 (AR1) was included to correct for temporal autocorrelation in the data. We expected that an increase in OD over the selection cycles will be *i*) stronger in AS than NS (i.e. significant effect of (*βγ*)_*j*_ ∗ *l*); *ii*) stronger in communities with high initial richness than in the ones with low richness (i.e. significant effect of (*αγ*)_*i*_ ∗ *l*); and that the overall gain in OD will be *iii*) stronger at high richness level in AS than in NS (i.e. significant effect of (*αβ*)_*ij*_). Our main hypothesis regarding an increase in selection efficiency with the increase in richness would be verified if *iv*) the increase in OD over the course of the experiment is stronger in AS than NS when richness increases (i.e. significant effect of the three-way interaction (*αβγ*)_*j*_ ∗ *l*). The analysis was done on the selected wells only in order to balance the dataset between AS and NS (one OD value per individual, cycle and selection treatment). The OD values were log10 transformed in order to meet the criteria for normality and homoscedasticity.

We detected a plate effect in the post-selection experiment: the OD of the control strains of two of the plates was lower than the observed OD for the other plates. We calculated the difference in OD and add this value to the measured OD of the two plates. Note that this correction did not significantly influence the results of the analysis. In order to compare what was observed during the experimental evolution and what was obtained from the post selection experiment, the data were analysed with the following linear mixed model:

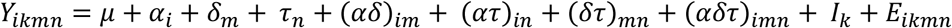

*Y*_*ikmn*_ is the OD of the individual of identity *k*, of initial richness level *i*, of history *m*, in dataset *n. µ* is the intercept, *α*_*i*_ is the effect of the initial richness level (qualitative: 1, 2, 4, 8, 16), *δ*_*m*_ is the effect of the history (qualitative: ancestor, evolved under AS, evolved under NS), *τ*_*n*_ is the effect the dataset (qualitative: experimental evolution, post-selection). The interaction effects between i) the initial richness level and the history (*αδ*)_*im*_; ii) the initial richness level and the dataset (*ατ*)_*in*_; iii) the history and the dataset (*δτ*)_*mn*_; iv) the initial richness level, the history and the dataset (*αδτ*)_*imn*_ were also included in the model. *I*_*k*_ is the random effect of the individual and *E*_*ikmn*_ is the residual error.

The description of the growth dynamics allowed to produce growth curves for each of the individual of the experiment under the three evolutionary histories (ancestor, evolved under AS, evolved under NS). We described the growth curves with segmented regressions which allowed to get the slopes of the different growth phases (four slopes) and the time of transition from one phase to the other (three breakpoints). From the growth curves, we also extracted the OD at 3.5 days (i.e. the targeted phenotype), the maximum OD and the time to reach the maximum OD. The obtained values for the three replicates of one individual were averaged. The data were analysed with the following linear model:

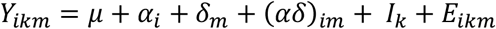

*Y*_*ikm*_ is the growth parameter of the individual of identity *k*, of initial richness level *i*, of history *m. µ* is the intercept, *α*_*i*_ is the effect of the initial richness level, *δ*_*m*_ is the effect of the history, (*αδ*)_*im*_ is the effect of the interaction between the initial richness level and the history. *I*_*k*_ is the random effect of the individual and *E*_*ikmn*_ is the residual error. A principal component analysis (PCA) including the ten growth parameters was performed and Euclidean distances between ancestors and evolved under AS and NS were computed based on the coordinates given by the PCA.

The same linear model as for the growth dynamic description experiment was used to analyse the number of metabolized substrates by the ancestors and evolved monocultures and communities (see Appendix 3 for details on the distinction between metabolized and non-metabolized substrates). The level of substrate use (i.e. the maximum OD reached on the different substrates) was analysed with the following linear mixed model:

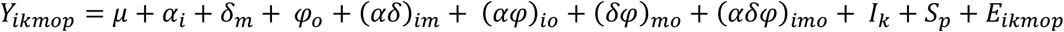

*Y*_*ikmop*_ is the OD of the individual of identity *k*, of initial richness level *i*, of history *m*, for substrate category *o* and substrate *p*. *µ* is the intercept, *α*_*i*_ is the effect of the initial richness level, *δ*_*m*_ is the effect of the history. *φ*_*o*_ is the effect of the substrate category. The interaction effects between i) the initial richness level and the history (*αδ*)_*im*_; ii) the initial richness level and the substrate category (*αφ*)_*io*_; iii) the history and the substrate category (*δφ*)_*mo*_; iv) the initial richness level, the history and the substrate category (*αδφ*)_*imo*_ were also included in the model. *I*_*k*_ is the random effect of the individual, *S*_*p*_ is the random effect of the substrate and *E*_*ikmnop*_ is the residual error. A PCA was conducted with the 31 substrates as variables.

All the analyses were performed with R version 3.6.3 with the following packages: nlme (Pinheiro et al. 2021) and lmerTest (Kuznetsova et al. 2017) for linear mixed models, car for type II analyses of variance (Fox and Weisberg 2019), emmeans for slope calculation (Lenth 2021), mclust for mixture models (Scrucca et al. 2016), segmented for segmented regressions (Muggeo 2008), FactoMineR for PCA (Lê et al. 2008).

## Results

### Artificial selection and initial richness level effects on mean OD

All selection cycles together, the OD in AS was significantly higher than this observed for NS (selection treatment: *χ*^2^=89; p_df=1_<2.2×10^−16^; Table 2): AS produced a gain in OD of 0.11, i.e. +16.4% as compared to NS. However, considering population mean in AS rather than the selected individuals only, the gain in OD in AS as compared to NS was +0.036, i.e. +5.3% (*χ*^2^=15; p_df=1_=1.1×10^−4^). There was also a significant effect of the initial richness level on OD (*χ*^2^=23, p_df=4_=1.5×10^−4^; Table 2). All selection cycles together, the OD tended to increase with the increase in the initial richness level (from 0.56±0.28 to 0.89±0.12) with significant differences between monocultures and levels 8 and 16.

**Table 2.**
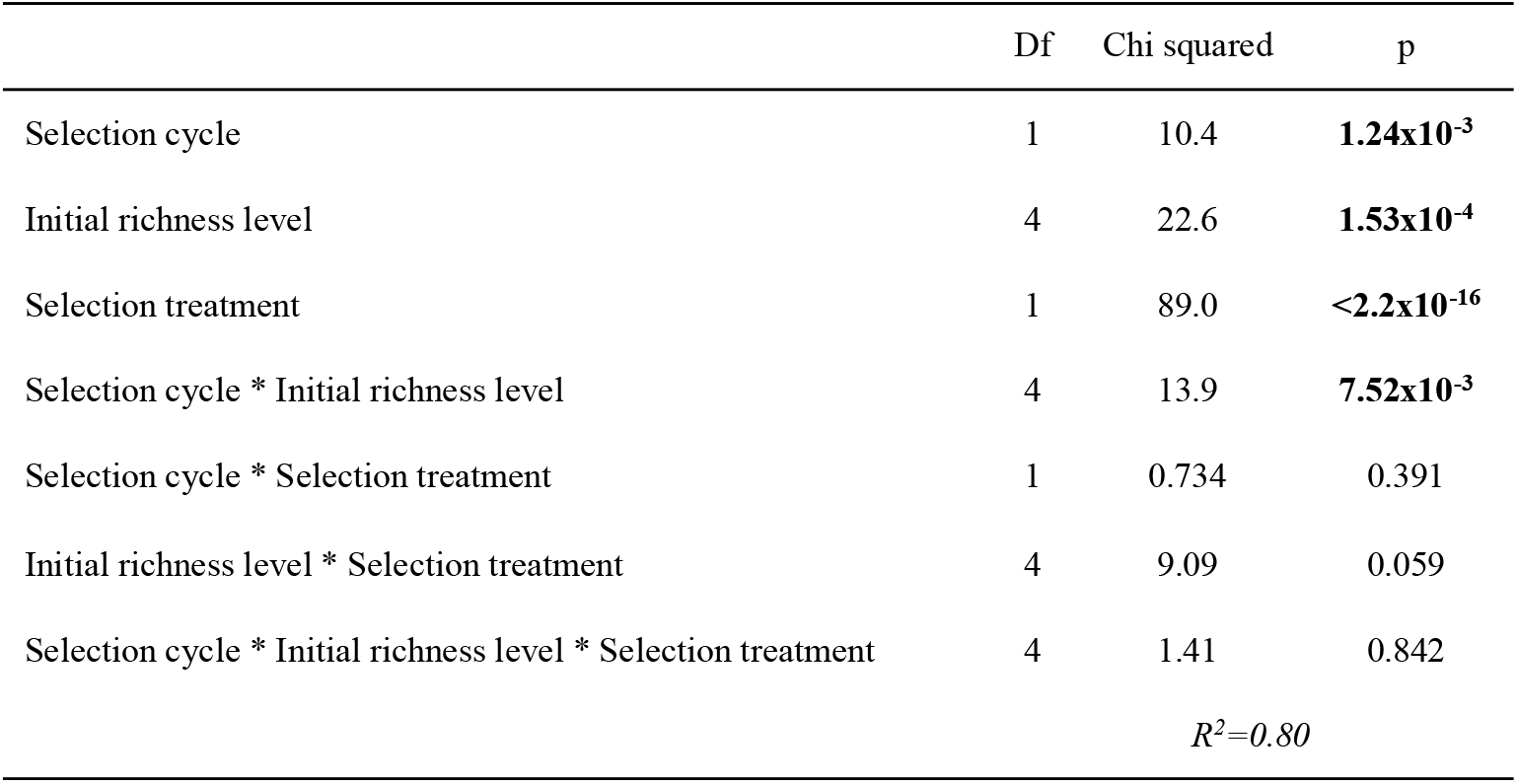
Deviance table of the covariance analysis (ANCOVA) of the optical density (OD) through experimental evolution. The effect of the selection cycle (from 1 to 40), the initial richness level (1, 2, 4, 8, 16), the selection treatment (artificial selection, no artificial selection) and their interactions on OD were estimated with a linear mixed model including the identity of the selection unit as a random effect factor and an autoregression structure. The conditional R^2^ is presented (i.e. variance explained by both fixed and random effect factors; the marginal R^2^ – fixed effect factors only – was 0.33).

|  | Df | Chi squared | p |
| --- | --- | --- | --- |
| Selection cycle | 1 | 10.4 | **$1.24 \times 10^{-3}$** |
| Initial richness level | 4 | 22.6 | **$1.53 \times 10^{-4}$** |
| Selection treatment | 1 | 89.0 | **$< 2.2 \times 10^{-16}$** |
| Selection cycle * Initial richness level | 4 | 13.9 | **$7.52 \times 10^{-3}$** |
| Selection cycle * Selection treatment | 1 | 0.734 | 0.391 |
| Initial richness level * Selection treatment | 4 | 9.09 | 0.059 |
| Selection cycle * Initial richness level * Selection treatment | 4 | 1.41 | 0.842 |
| | | | $R^2=0.80$ |

### OD of ancestors and evolved monocultures and communities

The OD of the monocultures and communities at the end of the experiment (i.e. evolved under AS or NS) differed from the OD of the monocultures and communities at the beginning of the experiment (i.e. ancestors). The OD of the ancestors was lower than the one of the evolved under NS which was lower than the one of the evolved under AS (0.58±0.23, 0.71±0.27 and 0.80±0.24 respectively; *χ*^2^=83; p_df=2_<2.2×10^−16^; Table S1). Thus, the artificially selected monocultures and communities were more productive than the ancestors and than the evolved under NS. This result did not depend on the initial richness level (initial richness level*history: *χ*^2^=3.0; p_df=8_=0.93; Table S1) and was consistent when considering either the OD retrieved from the experimental evolution or the OD measured in the post-selection experiment (Figure 1a and b; no effect of the dataset: *χ*^2^=1.2; p_df=1_=0.28; Table S1). However, when OD was measured in different growth conditions than those of the experimental evolution (i.e. in the system used to assess growth dynamics), the effect of the evolution depended on the dataset (*χ*^2^=329; p_df=4_<2.2×10^−16^) and the evolved under AS and NS showed a significantly lower OD than the ancestors (respectively 0.97±0.13, 0.95±0.16 and 1.1±0.22; Figure 1c) indicating that the abiotic environment influenced the expression of the phenotype under selection.

**Figure 1.**
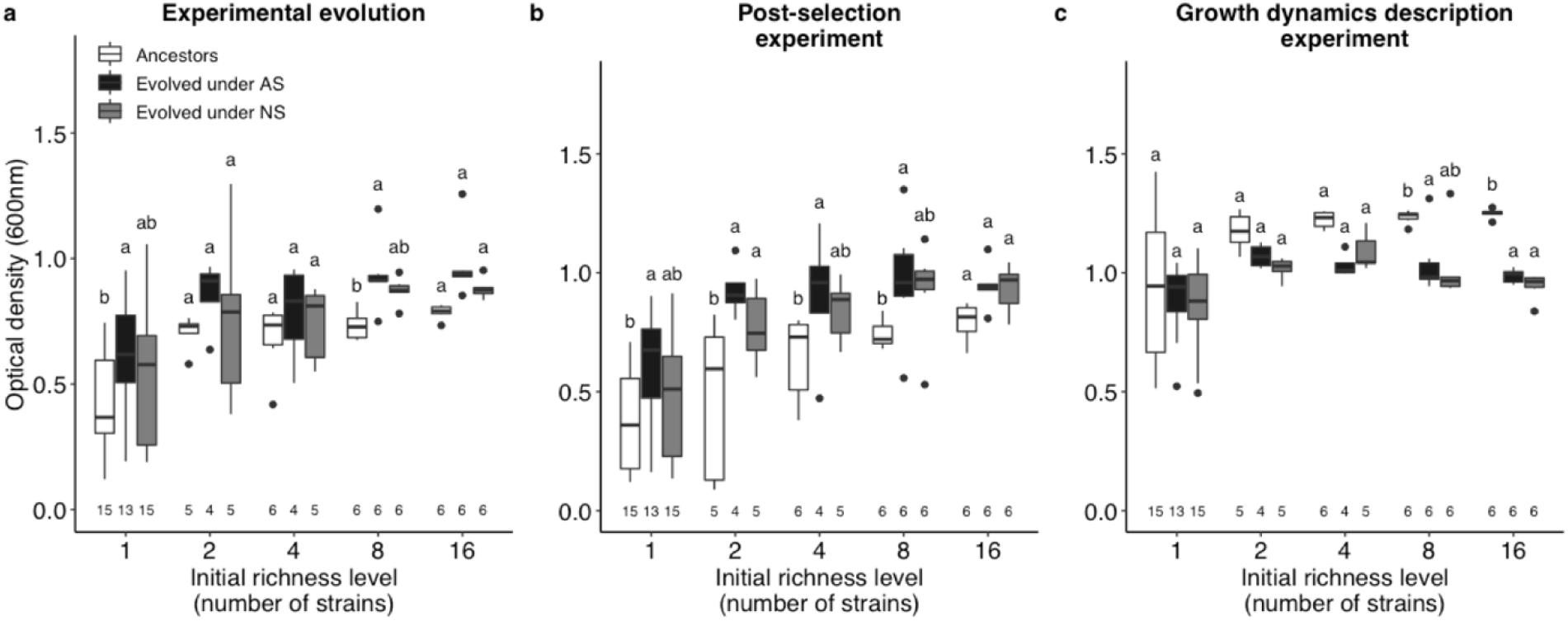
Optical density (OD) after 3.5 days of growth of ancestors and evolved monocultures and communities under artificial selection (AS) and no artificial selection (NS) depending on the initial richness level. a: OD measured during the experimental evolution experiment. The values of the ancestor corresponded to the values measured at cycle 1 (mean of AS and NS). The values of the evolved under AS and NS corresponded to the values measured at cycle 40. b: OD measured in the post-selection experiment (i.e. in the same experimental conditions). c: OD measured in the growth dynamics description experiment (i.e. in different experimental conditions). Each box represent the first quartile, the median and the third quartile for a given treatment, the end of the bars shows the minimal and maximal values within 1.5 times the interquartile range. The points outside of the boxes represent outliers. Sample sizes are given on the bottom of the graphs. Different letters represent significant differences between the histories within a richness level. White: ancestors; black: evolved under artificial selection; grey: evolved under no artificial selection.

### Artificial selection and initial richness level effects on OD change over time

There was no significant effect of the selection treatment on OD change over time (selection cycle*selection treatment: *χ*^2^=0.73; p_df=1_=0.39; Table 2). It indicated that, all richness levels together, the slope of the OD over the selection cycles was not different between AS and NS. On the contrary, the OD change over time was influenced by the initial richness level (selection cycle*initial richness level: *χ*^2^=14; p_df=4_=7.5×10^−3^; Table 2): OD tended to increase over time for the lowest (monocultures) and highest richness levels (eight and 16 strains) whereas it tended to decrease for intermediate richness levels (two and four strains; Figure 2).

**Figure 2.**
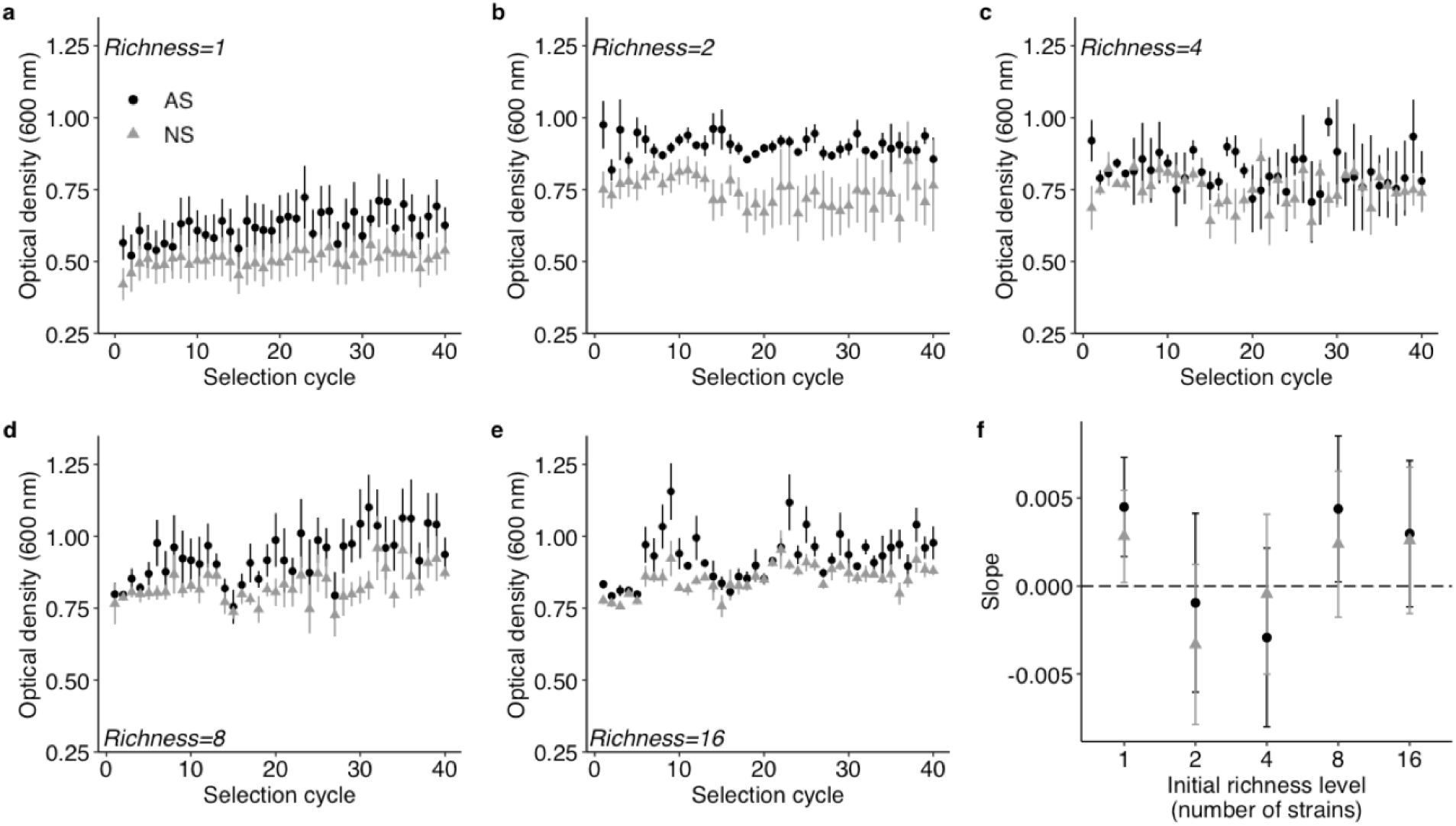
Changes in optical density (OD) over experimental evolution under artificial selection (AS) and no artificial selection (NS) depending on the initial richness level. a to e: The mean OD of the parents of the next population of selection units is represented by black circles for AS and grey triangles for NS for each initial richness level. Bars represent SE. Richness 1: n=13 in AS and 15 in NS (n=14 at cycle 33). Richness 2 and 4: n=4 in AS and 5 in NS. Richness 8 and 16: n=6 in AS and NS. The OD at cycle 0 was equal to 0.002 for all the treatments. f: The mean slopes of the regression lines predicted by a linear mixed model are presented in black circles for AS and grey triangles for NS for each initial richness level. Bars represent 95% CI.

### Artificial selection efficiency regarding the initial richness level

The outcome of artificial selection (AS) as compared to no artificial selection (NS) did not differ between the initial richness levels (selection cycle*selection treatment*initial richness level: *χ*^2^=1.4; p_df=4_=0.84; Table 2; Figure 2). It indicated that AS was not significantly effective whatever the initial richness level. However, there is still evidence for a possible influence of AS on OD change over time as compared to NS. Indeed, the slope of the OD over the selection cycles tended to be higher in AS than in NS in four richness levels over five (Figure 2f). Also, the differences in the slopes between AS and NS tended to be influenced by the initial richness level. Among the richness levels that showed an increase in OD over time, the highest differences in slopes between AS and NS occurred for level 8, where the slope in AS significantly differed from zero contrary to the slope in NS (4.4×10^−3^ and 2.4×10^− 3^ respectively), and for monocultures. On the contrary, the smallest difference occurred with an initial richness of 16 strains where both slopes were very similar (3.0×10^−3^ and 2.7×10^−3^ for AS and NS respectively; Figure 2f). It suggested that AS may have differentially influenced the increase in OD over time as compared to NS depending on the initial richness level. However, none of the richness levels responded enough to AS to observe significant differences. Interestingly, the correlation of the offspring-parent phenotype in AS was significantly higher in level 8 and monocultures than in level 16 (whereas it was not the case in NS; Figure S3), indicating a changing reliability of phenotype transmission with the change in the initial richness level.

### Artificial selection efficiency within the initial richness levels

Considering the detail of the response of OD through time for each individual or community within a richness level, we noticed that the variability of the response tended to decrease with the increase in initial richness (standard deviation of the mean slope in AS and NS together of 1.0×10^−2^ and 1.6×10^−3^ for monocultures and level 16 respectively; Figure S4). It indicated that the changes in OD over time were more similar between richer communities than between less rich ones or monocultures, where the responses to evolution were more contrasted. Furthermore, there was a high variability in the difference in slope between AS and NS within a richness level, especially at low richness levels (i.e. in monocultures and richness level 2). In monocultures, the two highest differences in slopes between AS and NS occurred for the two *Arthrobacter* strains (2.0×10^−2^ and 1.1×10^−2^ for *Arthrobacter* sp. BS2 and *Arthrobacter* sp. respectively). The two lowest differences between AS and NS occurred for two *Pseudomonas* strains (−2.4×10^−4^ and 8.0×10^−4^ for *Pseudomonas* sp. ADPe and *Pseudomonas knackmussii* DSM 6978 respectively; Figure S4). Thus, certain strains, and maybe genera, seemed to be more responsive to AS than others.

### Growth dynamics of ancestors and evolved monocultures and communities

Overall, the growth parameters of the ancestors and evolved differed more in communities than in monocultures (grouping of the ancestors on Figure 3b but not on 3a). However, considering the difference in growth parameters between ancestors and evolved individual by individual (i.e. looking at the distance between the ancestor and the corresponding evolved monoculture or community), there was no difference between monocultures and communities (mean Euclidean distance of 2.6±2.1 and 2.9±1.1 respectively). It highlighted the variability of the response in monocultures in which certain strains showed strong differences in growth parameters between ancestors and evolved whereas other showed no changes (Figure S5a and b). All richness levels together, the growth parameters tended to differ more between ancestors and evolved (Euclidean distance of 3.0±1.6 between ancestor and evolved under AS and of 3.3±1.6 between ancestor and evolved under NS) than between evolved under AS and evolved under NS (1.8±1.2). Indeed, the growth parameters changed in the same direction between evolved under AS and evolved under NS. There was an increase in the slope of the exponential growth phase as compared to the ancestors, which was significant for evolved under AS only (Figure S6a; Table S2), and a decrease in the time to reach the maximum OD (3,263 min±926, 3,404 min±986 and 4,416 min±534 for evolved under AS, evolved under NS and ancestors respectively; Figure S6b; Table S2). The difference in growth parameters between evolved under AS and evolved under NS tended to be lower in monocultures as compared to communities (Euclidean distance of 0.98±0.72 and 2.3±1.1 respectively), indicating a possible difference in the potential to phenotypic change under AS.

**Figure 3.**
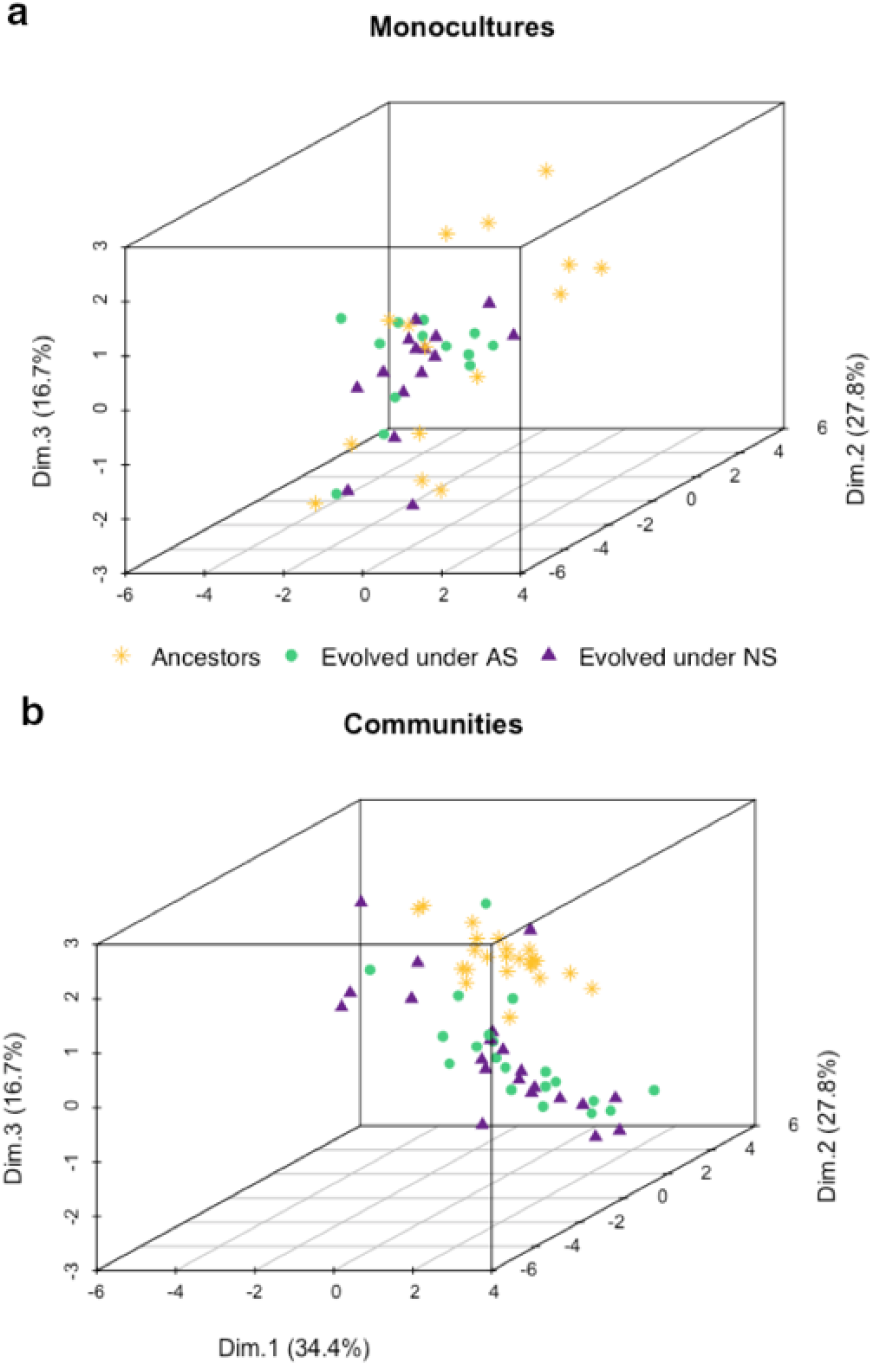
Principal component analysis of the growth parameters of ancestors and evolved monocultures (a) and communities (b). The ten growth parameters included in the analysis were retrieved from the description of growth curves obtained by repeated optical density (OD) measurements over 3.5 days. Those parameters were: the four slopes of the different growth phases, the three times associated to the transition from one phase to the other, the OD at 3.5 days, the maximum OD and the time to reach the maximum OD. The results obtained for monocultures and communities are presented separately for readability but were obtained from a unique analysis. Star: ancestors; circle: evolved under artificial selection; triangle: evolved under no artificial selection.

### Metabolism of ancestors and evolved monocultures and communities

The number of metabolized substrates increased with the increase of the initial richness level (it differed significantly between monocultures and the other richness levels; *χ*^2^=64; p_df=4_=5.0×10^−13^; Figure 4). It highlighted the existence of substrate use complementarity between the strains of the experiment. There was neither an effect of the history on the number of metabolized substrates (*χ*^2^=5.7; p_df=2_=0.06) nor an effect of the interaction between the history and the initial community richness (*χ*^2^=13; p_df=8_=0.11). It indicated that metabolic profiles were stable throughout the experimental evolution.

**Figure 4.**
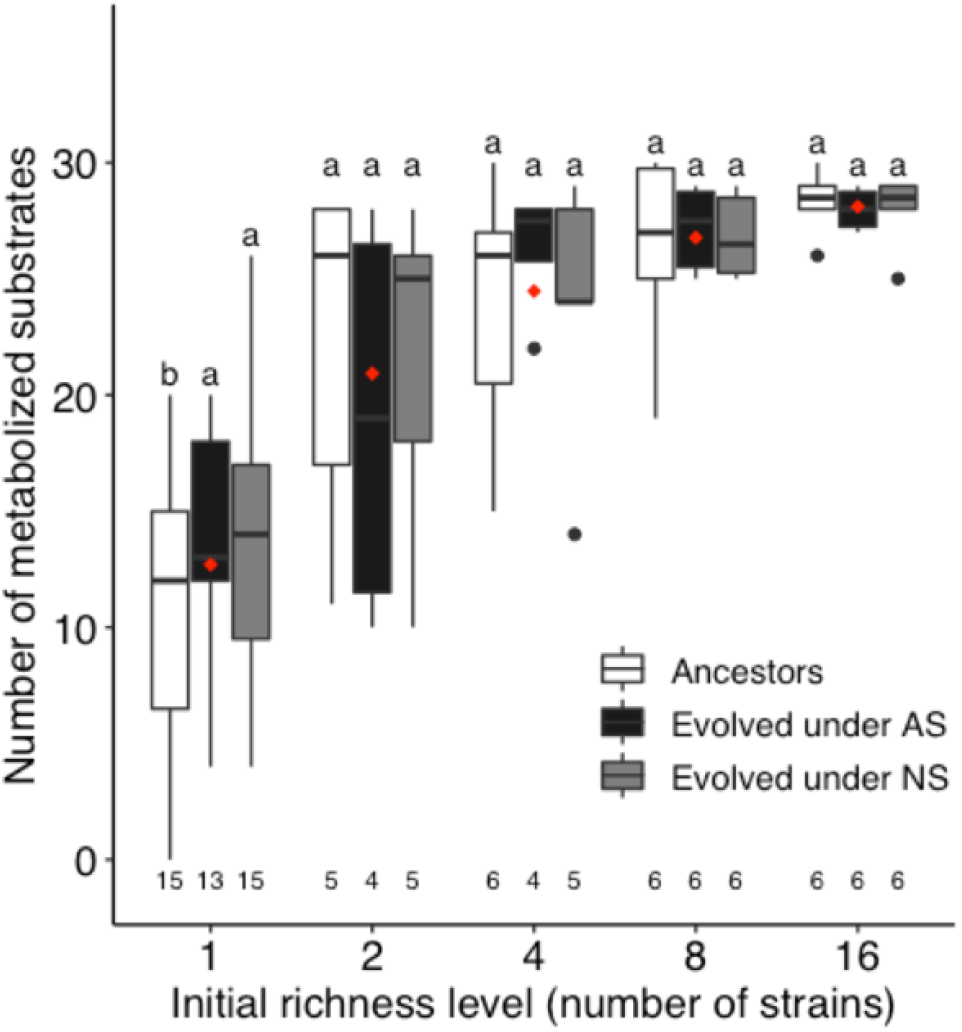
Number of metabolized substrates by ancestors and evolved monocultures and communities depending on the initial richness level. 31 substrates were tested in total. White: ancestors; black: evolved under artificial selection; grey: evolved under no artificial selection. Red diamonds represent the mean value for a given initial richness level. Each box represent the first quartile, the median and the third quartile for a given treatment, the end of the bars shows the minimal and maximal values within 1.5 times the interquartile range. The points outside of the boxes represent outliers. Sample sizes are given on the bottom of the graphs. Different letters represent significant differences between the histories within a richness level.

The OD reached on the 31 considered substrates was influenced by the initial richness level (*χ*^2^=43; p_df=4_=1.2×10^−8^; Table S3) and tended to be lower at lower richness levels (Figure 5a). There was an effect of the interaction between the initial richness level and the history (*χ*^2^=39; p_df=8_=6.1×10^−6^; Table S3). On the one side, the OD of the evolved tended to be higher than this of the ancestors for monocultures and richness levels 4 and 8. On the other side, the OD of the communities evolved under AS was lower than this of the communities evolved under NS at level 4 and lower than this of the ancestors at level 16 (Figure 5b). Thus, there was a trend to a gain or a loss in metabolic capabilities throughout evolution which depended on the initial richness level. The effect of the initial richness level on OD depended on the substrate category (*χ*^2^=127; p_df=20_<2.2×10^−16^; Table S3). Whereas the maximum OD tended to be achieved at level 4 for the phenolic compounds, the amines, the polymers and the amino acids, the OD tended to increase with the increase in richness for carbohydrates (maximum OD at level 8) and carboxylic acids (maximum OD at level 16; Figure 5c). It indicated that complementarity between the different strains occurred for certain substrates only.

**Figure 5.**
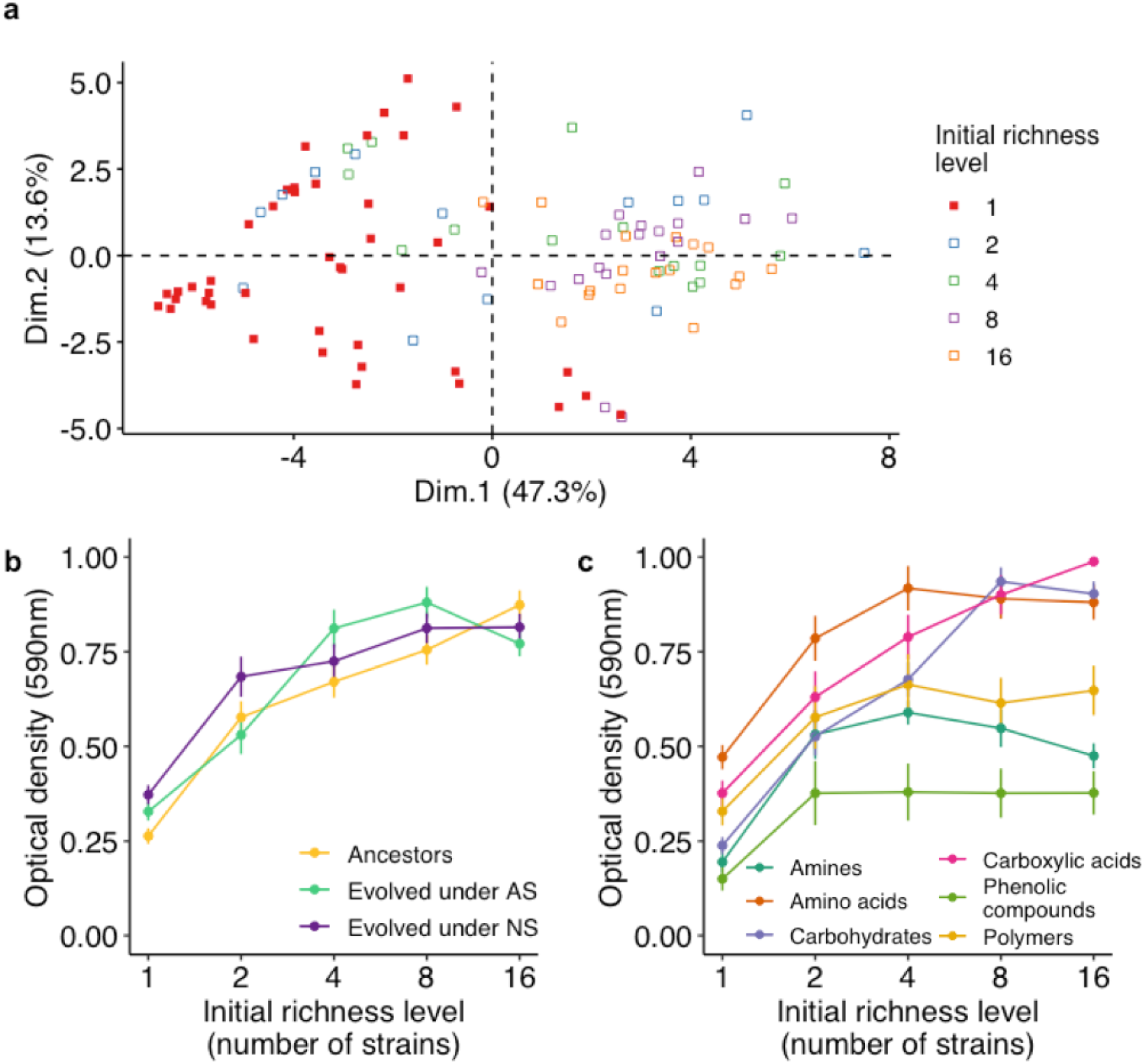
Substrate use depending on the initial richness level. a: Principal component analysis of the optical density measured on 31 carbon substrates for monocultures and communities. Ancestors, evolved under artificial selection and evolved under no artificial selection are all represented on the graph without distinction. The more a point is on the right of the graph, the more the corresponding strain or community reached a high OD on the tested substrates. b: Mean OD reached on the 31 tested substrates depending on the initial richness level and the history. Yellow: ancestors; green: evolved under artificial selection; violet: evolved under no artificial selection. c: Mean OD reached on each substrate category depending on the initial richness level. Bars represent SE. From lowest to highest value at level 1: phenolic compounds, amines, carbohydrates, polymers, carboxylic acids, amino acids.

## Discussion

Our results showed that artificial selection had an effect on the mean productivity of the bacterial communities and that community richness influenced both the mean productivity and productivity change over time (Table 2, Figure 2). Contrary to what was expected, there was no effect of the artificial selection on productivity change over time i.e. no increase in the artificially selected function. Previous studies showed significant changes over time in the selected function as compared to control treatments (Swenson et al. 2000a; Blouin et al. 2015; Chang et al. 2020). It is quite common however that artificial selection produces effects on the mean of a function rather than on the slope of function change versus time (Swenson et al. 2000b; Raynaud et al. 2019; Chang et al. 2020). The difficulty to observe a global trend in the change in a function under artificial selection is probably due to a lack of heritability, associated with the absence of stability in community structure (Chang et al. 2020; Jacquiod et al. 2021). A model developed by Xie et al. (2019) highlighted that the phenotypic variation in a community function is mainly due to non-heritable determinants, such as variation due to pipetting (when inoculating for the creation of a new generation) or function measurement noise for example. In accordance, in our experiment, the mean difference in OD between AS and NS was of 0.11 when considering the selected parents of the next cycle whereas it dropped to 0.036 when considering the population mean, revealing an important part of un-transmitted phenotypic variation. The absence of change in the slope could also be due to natural selection preventing artificial selection to be effective. Indeed, in artificial selection of communities, natural selection is also at stake within a selection unit, as observed in the NS treatment. In AS treatment, within-unit natural selection may overwhelm between-unit artificial selection, making the latter inefficient, as suggested in Wilson and Sober (1989) and Arora et al. (2020).

Species diversity could have multiple effects in artificial selection experiments. In our study, the initial richness level of the community influenced the selected function (productivity) and other potentially related ecosystem functions (i.e. growth dynamics, metabolic profile and level of substrate use, Figures 4 and 5) as often observed in diversity-functioning experiments (Bell et al. 2005; Gravel et al. 2011; Fetzer et al. 2015). Sequencing data indicated that there was dominance in the communities from the beginning of the experiment (Figure S7). Indeed, when the initial composition of a community included both *Escherichia coli* and *Pseudomonas* sp. ADP (ADP3 or ADPe) strains, almost no other species than these two was detectable. Similar outcomes were observed in previous studies. In Goldford et al. (2018), despite the various origins of the twelve studied communities and their high initial richness level (from 110 to 1,290 exact sequence variants (ESV)), all of the communities converged to the same composition at the family level, i.e. Enterobacteriaceae and Pseudomonadaceae (from 4 to 17 ESV) after 12 serial transfers. This family-level composition was also observed in the study of Scheuerl et al., (2020) in all of the 64 studied communities after a five-month experimental evolution. Enterobacteriaceae and Pseudomonadaceae strains retrieved from the evolved communities in Goldford et al. (2018) were all able to grow on the metabolic by-products of all other strains of the community, indicating that cross-feeding was at stake in these communities which may also have occurred in our experiment. Based on the sequencing data, the *Escherichia-Pseudomonas* co-dominance structure occurred twice at level 2, three times at level 4, five times at level 8 and six times at level 16 (in both selection treatments). Thus, the observed positive diversity-functioning relationship could be due to the increased probability of selecting the cross-feeding community members by increasing the initial richness level (i.e. a sampling effect of the complementarity effect).

The effect of species richness in an artificial selection experiment can also occur through an interaction with the evolutionary dynamics or with the selection treatments. In our study, the initial richness level did not significantly influence the effect of the selection treatment (initial richness level*selection treatment, p_df=4_=0.059) as the mean productivity was always (and similarly) higher in AS than in NS whatever the community richness. However, community richness influenced the evolutionary dynamics (initial richness level*selection cycle, p_df=4_=7.52×10^−3^, Table 2). The sign of the effect depended on the initial richness level but also on the considered community within a richness level; it suggested an influence of community composition on the community evolutionary trajectory. Thus, as presented in the literature, our results suggest that there is an interplay between community ecology and community evolution (Johnson and Stinchcombe 2007; O’Brien et al. 2013) and indicate that the effect of community diversity could change with the timescale at which community function is considered. Species richness could also affect the way artificial selection influences the evolutionary dynamics (i.e. the efficiency of artificial selection, identified as the three-way interaction between the initial richness level, the selection cycle and the selection treatment in Materials and Methods section). We hypothesized that artificial selection efficiency would increase with the initial richness level through an increase in the sources of variations (species composition, intra and interspecific interactions…) and hence an increase in the existing solutions to reach the targeted phenotype. However, there was no significant difference in the slopes of the OD response versus time between AS and NS depending on community richness in a linear model (selection cycle*initial richness level*selection treatment, p_df=4_=0.842, Table 2). Nevertheless, there was still evidence that the initial richness level could influence artificial selection efficiency as the difference in OD change over time between AS and NS tended to be non-linearly affected by an increase in community richness (Figure 2f). Moreover, we noticed that the correlation between parent and offspring OD depended on the initial richness level and that it also responded idiosyncratically to an increase in richness (Figure S3). Previous modelling approaches highlighted that in artificial selection of communities, a fine balance between variation and heritability must be achieved (Penn 2003). Based on our results, we suggest that the search for this equilibrium could occur through the modulation of community diversity. But, in addition to the initial species richness, divergence between replicates within a richness level has to be considered to understand the effect of the initial diversity on the efficiency of artificial selection.

Increasing the initial richness level decreased the between-community variation within a richness level. This especially came with the design of our experiment but could also occur when working on natural communities. In our study, we built the different communities from an initial pool of 18 strains. As a consequence, whereas none of the communities of level 2 included strains in common, seven strains were found in the six communities of level 16 (and at least one *Escherichia* and *Pseudomonas* strain as discussed above; Table S4). It is well-known that community composition has an effect on community functioning as, for a given richness level, a panel of community phenotypes can be observed depending on community composition (Bell et al. 2005; Fetzer et al. 2015). Between- community differences in composition could also be potential levers for artificial selection. In a recent study, Sánchez et al. (2021) proposed that an efficient directed evolution of microbial community would occur through a good exploration of the “ecological structure-function landscape”. In order to explore more solutions to reach the targeted community phenotype (the “function” component of the landscape), multiple communities varying for their composition (the “structure” component of the landscape) have to be considered. In this idea, the first step of a directed evolution experiment would be to create a library of communities varying for their composition (Sánchez et al. 2021). In the light of our results, we suggest that the differences in community composition in the initial pool of selection units must be sharp enough (e.g. family-level differences) to avoid resemblance in community dynamics that would reduce the exploration of multiple evolutionary trajectories. A first solution could be to start from an initial pool of species several times higher than the number of species in the highest richness level (e.g. 64 instead of 18 species to build six replicates of the 16-species level that deeply differ in their composition). Another way to ensure sufficient compositional variability is the maintenance of multiple lineages over the experiment (Blouin et al. 2015; Jacquiod et al. 2021). In a recent study, (Chang et al. 2020) started from a pool of 12 communities which were replicated seven or eight times each for a total of 92 communities. A selection cycle occurred through the selection of the 23 best performing communities among the 92 and, after six selection cycles, all of the communities stemmed from an unique parental community (Chang et al. 2020). The single lineage increased the probability that the gain in the targeted function under AS was due to the elimination of the less performing communities but not to an increase in the function itself. Applying artificial selection within several independent lineages would prevent the results to be due to ecological sorting (i.e. a simple identification of the best performing communities in an initial pool) and enhance the probability of finding communities that are responsive to the selection.

In the present study, we showed that the diversity of the communities could play a role in the artificial selection procedures. Community richness had an effect on the selected property and influenced the community evolutionary dynamics. Also, we found evidence that it could impact the efficiency of artificial selection, but the trade-off between increasing richness and maintaining variability in composition makes the effect of the initial richness non-linear. Indeed, one of the limitations that can occur when increasing initial community richness from a limited pool of species is the convergence in community composition that may reduce between-community variations for artificial selection to act upon. Once this limitation is avoided, we suggest that applying artificial selection on community varying for their diversity could allow to explore multiple variability/heritability. Protocol optimization is still needed for artificial selection of microbial communities to be efficient and, multiple lines of improvement have already been highlighted by recent modelling approaches and experimental studies. Further studies will be needed to disentangle the links between community ecological dynamics and community evolutionary trajectory, which will open the way for effective microbial community and microbiome engineering.

## Acknowledgments

We thank David Bru and Jérémie Béguet from the EMFEED team, INRAE Dijon, for their advice on molecular biology analyses. This work was supported by grants from the INRAE, AGROECOSYS- TEM department, “Pari scientifique” program, ref 6503.

## Author contributions

A.S., M.B. and T.R. designed the study. M.D. and T.R performed the experiments. A.S. and T.R. analyzed the data. T.R. wrote the paper with substantial contributions of A.S. and M.B.

## Data sharing plans

All the data and codes used during the study will be available from the corresponding author on reasonable request.

