## Supplemental Figures and Tables for "Community diversity determines the evolution of synthetic bacterial communities under artificial selection"

**Supplementary material**

### Appendix 1: Community composition analysis

*Molecular biology*

In order to perform DNA extractions, monocultures and communities from cycles 1 (i.e. initial inocula after 84 h of growth) and 40 and communities from intermediate cycles 14 and 27 were revived from the glycerol stocks in the growth conditions of the experiment (i.e. deep-well plates, 28°C, no shaking, 84 h, the culture medium volume was increased to 1.8 ml following the instructions of the DNA extraction kit manufacturer; DNeasy UltraClean 96 Microbial Kit, Qiagen, Netherlands). After 84 h of growth, the plates were centrifuged for 24 min at 2,250 RCF, the supernatant was discarded, the plates were centrifuged again for 8 min at 2,250 RCF and the remaining supernatant was discarded. The obtained cell pellets were stored at -20°C. DNA extractions were then conducted following the kit manufacturer's instructions. The extracted DNA was quantified (Quant-iT dsDNA Assay Kit high sensitivity, Invitrogen, USA) and each sample was diluted to 0.5 ng. $\mu L^{-1}$. A mock community (mock 1) was created by adding an equivalent volume of extracted DNA (0.5 ng. $\mu L^{-1}$) of each of the 18 strains and included into the samples to be analysed as well as another mock community (mock 2) with other strains than those of the experimentation. We first performed 16S rRNA gene amplification (V3-V4 region) using the PCR primers Pro341F/Pro805R (carrying 5' tails for Illumina sequencing) at a final concentration of 0.25 µM for 1 ng of DNA. The cycling conditions were: 98 °C (3 min), 25 amplification cycles at 98 °C (30 s), 55 °C (30 s), 72 °C (30 s) and a final step at 72 °C (10 min). Then we performed a second PCR to allow the barcoding of the PCR products with NGS primers at a final concentration of 1 µM using the same cycling conditions but reducing the number of amplification cycles to eight. The obtained PCR products were normalized (SequalPrep Normalization Plate Kit, Applied Biosystems, USA), pooled and sequenced on Illumina MiSeq (2x250 bp; Genoscreen, France). We then performed *gyrB* gene (coding for the subunit B of DNA gyrase) amplification using PCR primers that were specifically designed for the 18 strains of the experiment: gyrB55F (5'-GTNMGHAARCGBCCNGS-3') and gyrB347R (5'-TGHARRCCRCCVGANAC-3') carrying the 5' tails for Illumina sequencing. We designed these primers to prevent nonspecific amplification of *parE* gene (coding for the subunit B of DNA topoisomerase IV) which shows high identity levels with *gyrB* as noticed by *in silico* testing of pre-existing *gyrB* primers. The primers were used in a final concentration of 2 µM for 1 ng of DNA. The cycling conditions were: 98°C (4min), 35 amplification cycles at 98 °C (45 s), 58 °C (1 min), 72 °C (1 min) and a final step at 72 °C (10 min). The obtained PCR products were purified (Pronex Size-Selective Purification System, Promega, USA) following the manufacturer's protocol to remove the unfixed PCR primers (approximate size cut-off of 250 base pairs) as they were degenerate. The next steps were identical to those of 16S rRNA gene: second PCR to attach the barcodes, normalization, pooling and Illumina MiSeq sequencing.

*DNA database construction*

As there is no available full length *gyrB* database, we built our own *gyrB* database. We isolated the forward and reverse sequences of isolated ancestral strains from the rest of the sequences obtained on the Illumina MiSeq run. We then assembled the forward and reverse sequences (PEAR; Zhang et al. 2014) and discarded unassembled sequences. Sequences smaller than 100 base pairs were discarded as well as chimeras (which were detected *de novo*). OTU clustering was done at the 94% identity level (VSEARCH; Rognes et al. 2016). An OTU table was built and the sequences of the dominant OTUs in each sample were aligned against GenBank database using BLAST (Altschul et al. 1990) to check for the strain and the gene identity before being added to the database. Sequences that did not correspond to *gyrB* were not included in the database except one sequence found for Arthrobacter sp. BS2 which showed 94% identity with a gene whose product is a topoisomerase IV (which is a product of *parE* gene) whereas there was no sequence that corresponded to *gyrB*. Each sequence that was included in the database was given an identifier and the corresponding taxonomy was reported in another file. We ran again the process starting from the same set of forward and reverse sequences, setting the minimal sequence length to 100 and adding a reference-based chimera check step with our database as reference. Once OTU clustering was done (94% identity) and the OTU table was built, the taxonomy was assigned using our reference database: each sequence was aligned against the reference sequences and the highest identity level was kept to assign the corresponding taxonomy provided that the alignment length was higher than 250 bp. The obtained sequences for each sample were added to the database (after checking for strain and gene identity) as well as sequences of the strains of the mock 2. There were four strains for which no sequence was retrieved with this procedure (*Aminobacter aminovorans* SR38, *Arthrobacter* sp, *Microbacterium* sp. C448, *Pseudopedobacter saltans* DSM 12145), *gyrB* sequences were collected from GenBank or personal data and added to the database. For sake of consistency, we followed the same procedure for the construction of the 16S rRNA gene database (minimal sequence length: 350 bp, OTU clustering at 94% identity, minimal alignment length: 350 bp).

*Identity threshold for OTU clustering*

Three replicates of two mock communities were sequenced and their analysis was used to choose the identity level threshold for OTU clustering by maximizing the number of detected strains (based on the taxonomy) while minimizing the number of OTUs. The first mock community (mock 1) was a combination of the 18 strains of the experiment belonging to 12 genera and 15 or 16 species. The second mock community (mock 2) was a mix of 39 strains (other than those of the experiment) belonging to 17 genera and 27 species. We tested identity level thresholds for OTU clustering ranging from 0.92 to 0.97. On *gyrB* sequences, the highest number of detected strains on mock 1 was 11 and was obtained with an identity level of 0.95; it corresponded to 56 OTUs (at 0.94 identity level we detected 10 strains and 29 OTUs and at 0.96 we detected 9 strains and 98 OTUs). On mock 2, the 0.95 identity level allowed to detect 24 strains and 130 OTUs. On 16S sequences, 11 strains were detected in mock 1 whatever the identity level with an OTU number varying between 22 and 97. On mock 2, the highest number of detected strains was 25 and was obtained with both 0.94 and 0.95 identity levels but with 82 and 101 OTUs respectively. We thus chose the 0.94 threshold (which corresponded to 36 OTUs on mock 1 against 32 with the 0.95 threshold).

*Bioinformatics analyses*

The forward and reverse sequences obtained from Illumina MiSeq sequencing were assembled using PEAR (Zhang et al. 2014) and unassembled sequences were discarded as well as sequences smaller than 200 bp for *gyrB* and 350 bp for 16S rRNA gene. OTU picking was conducted against the reference databases that we constructed. First, chimeras were eliminated with *de novo* and reference based chimera detection, then, OTU clustering was done using VSEARCH (Rognes et al. 2016) at identity thresholds of 0.95 and 0.94 for *gyrB* and 16S respectively (based on the results we obtained on the mock communities). One representative sequence per cluster was kept based on the highest base pair number. The taxonomy was assigned to each sequence based on the highest identity level with the sequences of our reference database and with a condition on the length of the alignment (> 200 and >350 bp for *gyrB* and 16S respectively). An OTU table was then constructed. Based on the expected community composition, we noticed the presence of unexpected OTUs with very low counts in several samples. This could be due to critical mistags i.e. a sequencing barcode is associated with the wrong sequence and this can not be detected as it corresponds to an existing barcode combination (which is highly probable as we used almost all possible barcodes combinations; Esling et al. 2015). Very low counts of an unexpected OTU can also be due to cross-talk, i.e. a read is assigned to the wrong sample at the demultiplexing time. Contrary to critical mistags, this can be corrected *a posteriori*. We estimated *de novo* the cross-talk rate for each OTU using the UNCROSS2 algorithm (Edgar 2018). For each OTU count in each sample, this algorithm gives a score *t* which is close to 0 when there is no indication for cross-talk and close to 1 when cross-talk is detected. When *t* was higher than 0.1 (i.e. the threshold recommended by Edgar 2018), the corresponding count was set to 0. The main OTU was identified for each strain and the corresponding corrected counts were included in the final OTU tables. From these corrected OTU tables for 16S rRNA gene and *gyrB*, we built one presence/absence OTU table based on the condition that one strain was present if the corresponding OTU counts was higher than 100 and zero for 16S and *gyrB* respectively. The final OTU tables contained only the counts of the strains coded as "present".

### Appendix 2: Contaminant management

After the experimental evolution, the experimentation was conducted in the following order starting from the same glycerol stocks: growth profile study, metabolism study, DNA extractions and PCR, post-selection experiment. Therefore, when a contamination occurred in a sample during the experimental evolution, it was detectable in all of the datasets and we removed the corresponding sample from the analyses. When the indication for possible contamination involved only one of the datasets, we considered that it was not sufficient to remove the corresponding sample from the analyses (see Figure S1 for more details). When a sample was contaminated in a given selection treatment (i.e. AS or NS), we removed this sample from the datasets for the affected selection treatment and kept the corresponding sample in the other selection treatment. When both selection treatments were affected, we removed the samples and the corresponding ancestor in the datasets. It resulted in the removal of 16.7% of the samples in the experimental evolution dataset, distributed as follows: artificial selection: 21.4%; no artificial selection: 11.9%; richness 1: 22.2%; richness 2: 25%; richness 4: 25%; richness 8: 0%; richness 16: 0%. In the growth profile dataset, metabolism dataset and post-selection dataset, it resulted in the removal of 14.3% of the samples distributed as follows: artificial selection: 21.4%; no artificial selection: 11.9%; ancestors: 9.5%; richness 1: 20.4%; richness 2: 22.2%; richness 4: 16.7%; richness 8: 0%; richness 16: 0%.

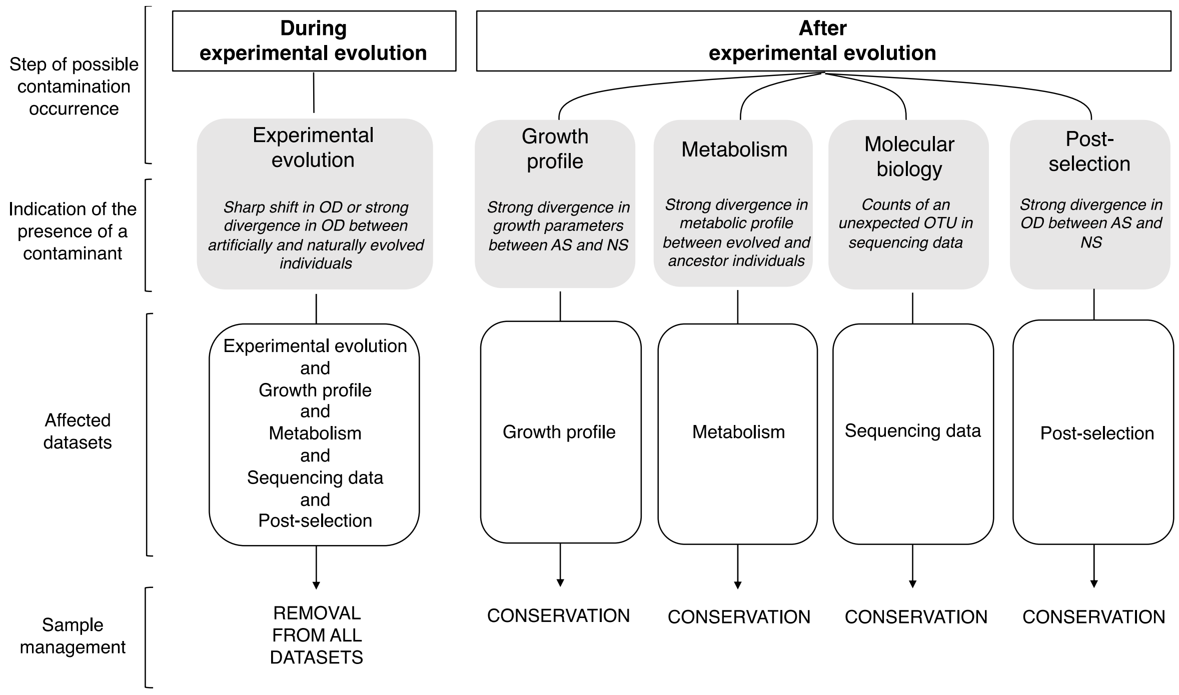


**Figure S1: Contaminant management in the datasets.** There were five different steps in the experimentation, each of which corresponding to a distinct dataset. The different steps are presented from left to right in the order in which they were conducted. The indication of the presence of a contaminant in a sample was defined as a deviation from the global trend observed for a given dataset (e.g. in metabolism dataset, the metabolic profile is overall very similar between ancestors and evolved individuals, a shift in metabolic profile thus suggested a possible contamination). If a contamination occurred during the experimental evolution, it affected all of the steps of the experimentation and thus, there were indications of the presence of a contaminant in all of the datasets. In this case, we considered that the amount of information was sufficient to remove the sample from the analyses with low error probability. On the contrary, when there was an indication of the presence of a contaminant in only one of the steps (which did not occur in the experimental evolution dataset), we considered that the amount of information was not sufficient enough to remove the sample. Another possibility was the contamination of the glycerol stocks from which the different steps were conducted, in this case we would have indications for the presence of a contaminants in at least two steps in a row which was not observed. OD: optical density; OTU: operational taxonomic unit.

**Appendix 3: Metabolism data analysis**

To analyse the metabolism data, we first calculated the difference between the OD produced from one substrate and the mean OD of the control wells (containing water) for each individual at each studied time (negative values were set to zero). Then, for each substrate, we determined a threshold value from which the OD was sufficiently different from zero to consider the substrate as metabolized. To determine this threshold value, we identified two populations in the values differing from zero using a mixture model, these two populations corresponded to a first group of values of low variance and high density (non-metabolized) and to a second group of values with a higher variance and a lower density (metabolized). The OD of the sample separating the two populations was taken as the threshold value (Figure S1a). For six substrates over the 31, the values had to be divided into three populations in order to distinguish the high-density low-variance population (Figure S1b). A substrate was considered as metabolized if the measured OD overpassed the threshold value for this substrate at least for the two last measurements in time and for the three replicates. For the metabolized substrates, we extracted the maximum OD and then described the metabolization dynamic by linear regression or linear segmented regression. For each sample, we ran regressions with zero to two breakpoints and kept the slope estimations of the regression showing the highest $R^2$. In order to compare the slopes whatever the regression used, we kept the maximum slope value for each sample. The maximum slope and maximum OD were averaged for the three replicates of an individual. These two indicators were highly correlated ($\rho$=0.96, $p<2.2\times10^{-16}$) therefore the analyses were conducted on the maximum OD only.

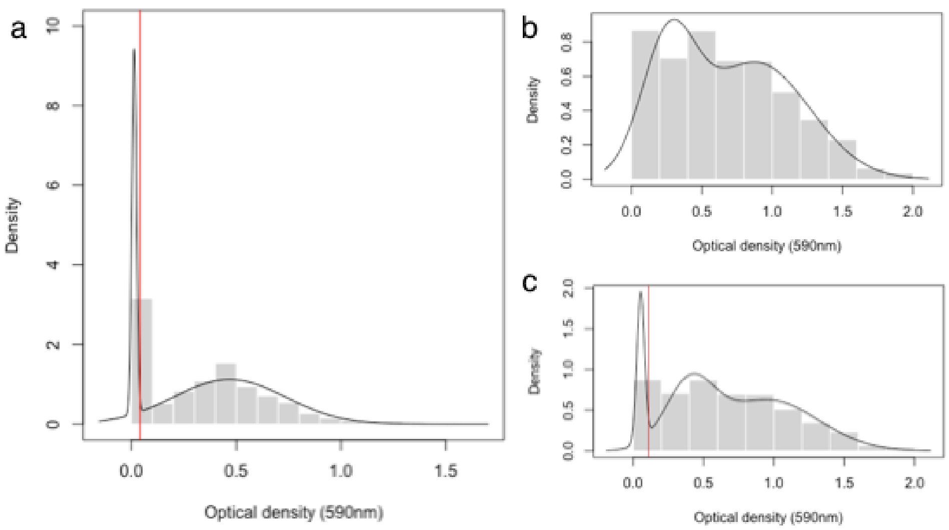


**Figure S2 Threshold value determination for metabolism data analysis.** For each of the 31 substrates studied in the experiment, we determined a threshold value of optical density (OD) from which we considered that a substrate was metabolized as compared to the OD measured on a control containing water. **a:** For β-methyl-D-glucoside, as for 24 other substrates, a mixture model with two gaussian distributions allowed to distinguish between a first population of value of high density and low variance (i.e. samples in which the substrate was not metabolized) and a second population of low density and high variance (i.e. samples in which the substrate was metabolized). In these cases, the threshold value (represented by a vertical red line) was the OD of the sample separating the two populations. **b:** For pyruvic acid methyl ester, as for five other substrates, a mixture model with two gaussian distributions did not allow to identify the population of values of high density and low variance. **c:** In these cases, fitting a mixture model with three gaussian distributions was necessary. The threshold value was the OD of the sample separating the first two populations.

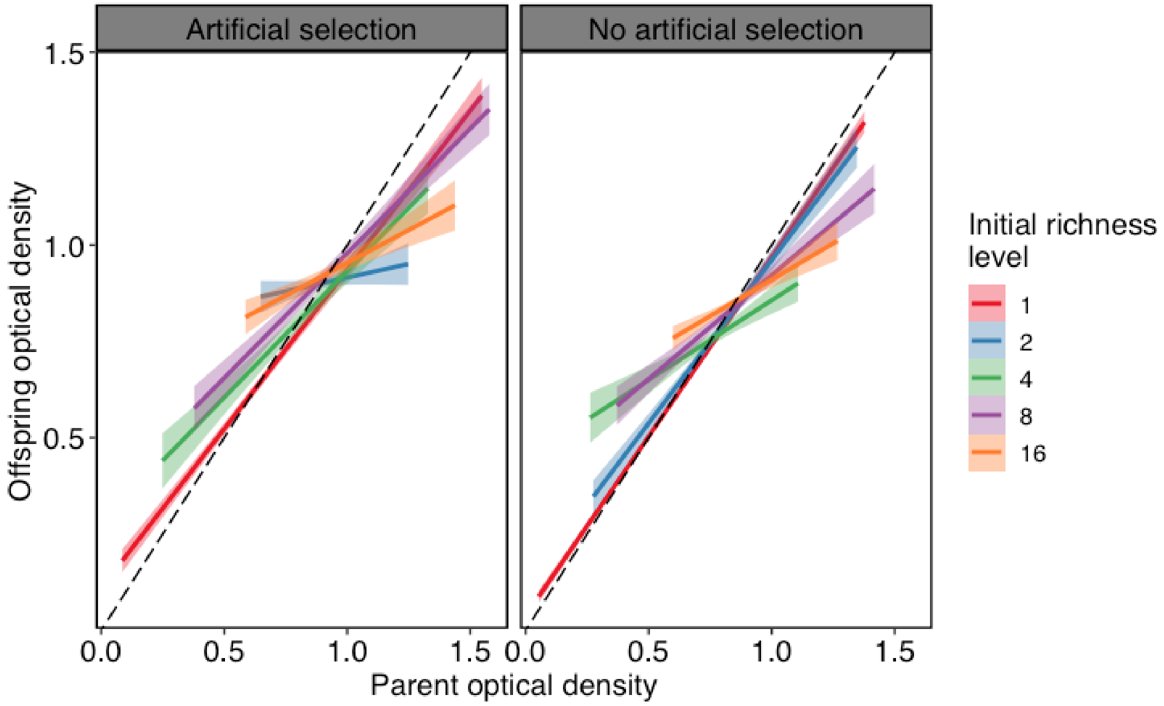


**Figure S3 Correlation between offspring and parent phenotype.** The optical density of the offspring (i.e. OD of the selected well at selection cycle n+1) is presented against the optical density of the parents (i.e. OD of the selected well at selection cycle n) for each initial richness level within each selection method (left: artificial selection, right: no artificial selection). The theoretical regression line for a correlation value of 1 is represented by a black dashed-line. There was a significant effect of the interaction parent optical density*initial richness level*selection method on the offspring optical density ($\chi^2$=28; $p_{df=4}$=1.2x$10^{-5}$).

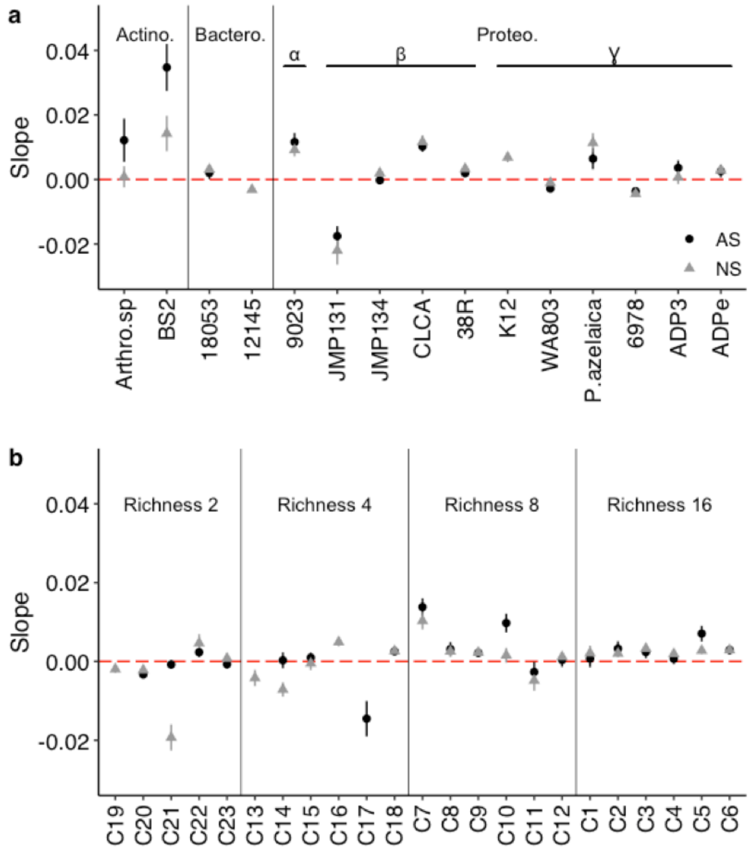


**Figure S4 Changes in optical density (OD) over the experimental evolution under artificial selection (AS) and no artificial selection (NS) for each monoculture (a) and community (b) of the experiment.** The slopes of the regression lines predicted by a linear mixed model (with the identity of the individual nested into the selection method as a random effect factor on the slope and the intercept) are presented in black circles for AS and grey triangles for NS for each initial richness level. Bars represent SE. Actino.: Actinobacteria, Bactero.: Bacteroidetes, Proteo.:Proteobacteria. $\alpha$: $\alpha$-Proteobacteria, β: β-Proteobacteria, ɣ: ɣ-Proteobacteria.

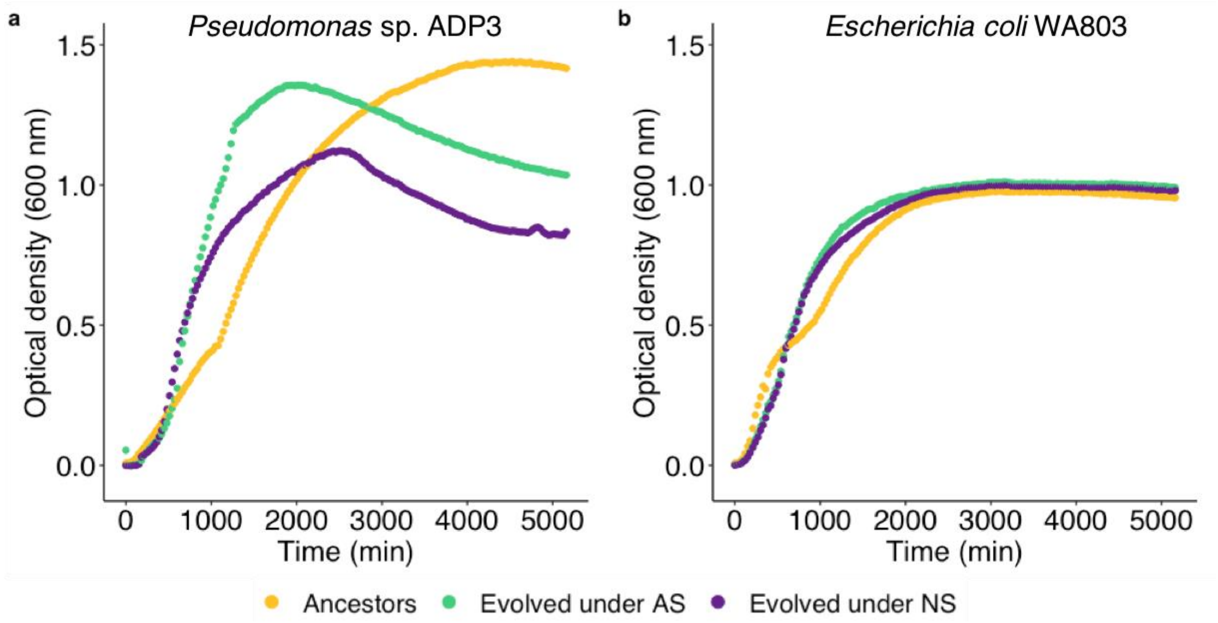


204

205

206 **Figure S5 Growth curves of two monocultures depending on their evolutionary history.** Each point
207 corresponds to the mean optical density of three independent measurements. Yellow: ancestors, green:
208 evolved under artificial selection, violet: evolved under no artificial selection.

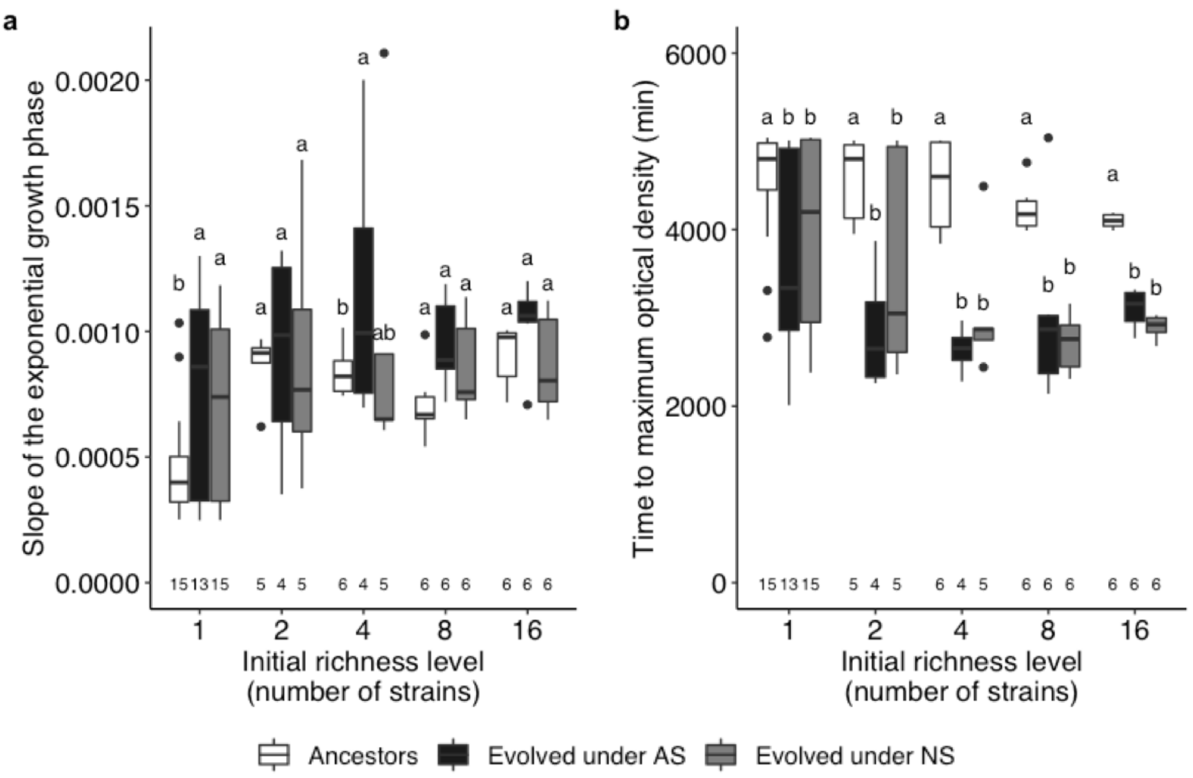


**Figure S6 Growth parameters of the ancestors and evolved monocultures and communities under artificial and no artificial selection depending on the initial richness level. a:** Slope of the exponential growth phase. **b:** Time to reach the maximum OD. Each box represents the first quartile, the median and the third quartile for a given treatment, the end of the bars shows the minimal and maximal values within 1.5 times the interquartile range. The points outside of the boxes represent outliers. Sample sizes are given on the bottom of the graphs. Different letters represent significant differences between the levels of history within a richness level. White: ancestors; black: evolved under artificial selection; grey: evolved under no artificial selection.

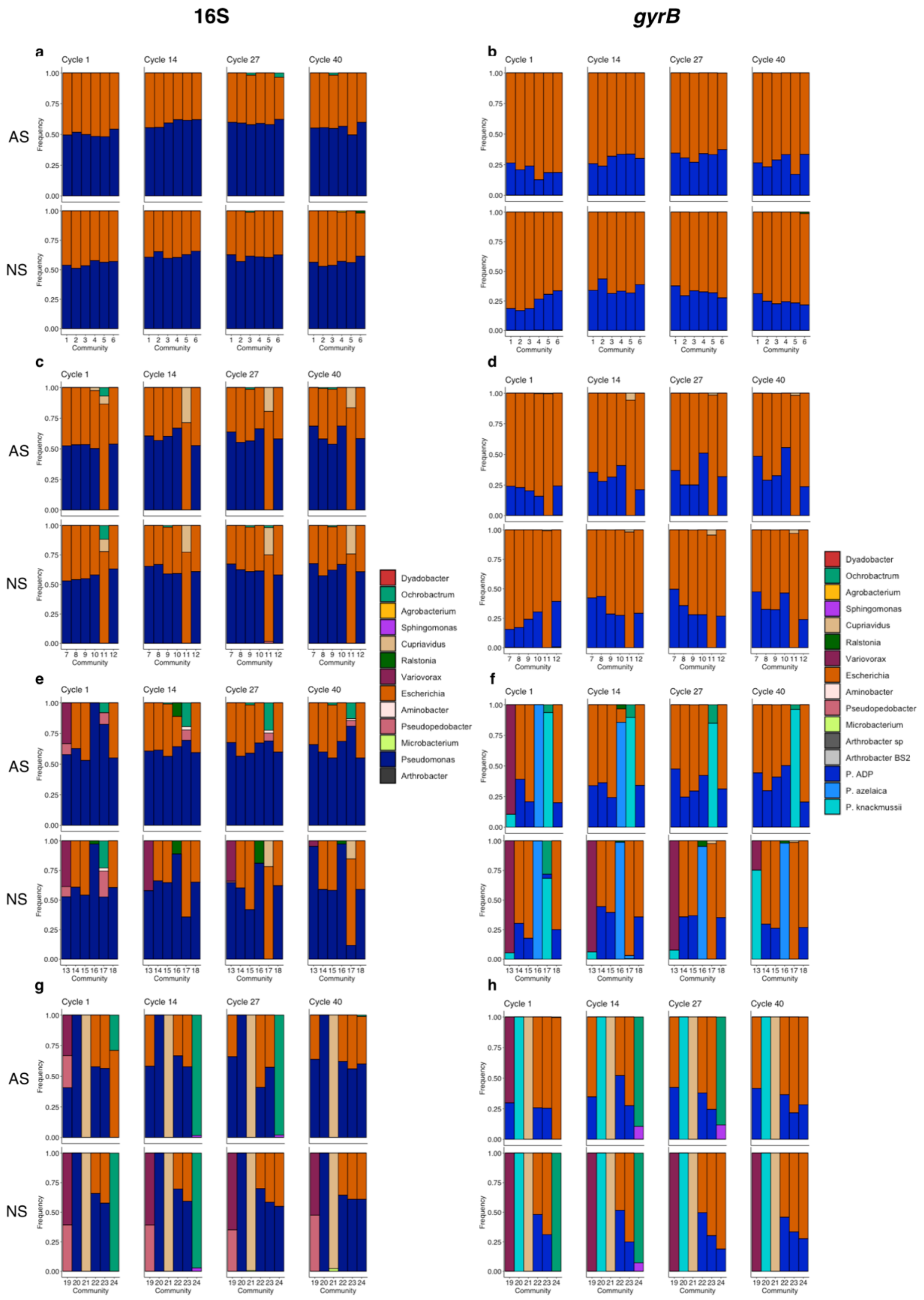


**Figure S7 Community composition over the selection cycles.** The composition of the communities was analyzed by 16S RNA gene sequencing (a, c, e, g) and by *gyrB* gene sequencing (b, d, f, h) at cycles 1, 14, 27 and 40 (from left to right) for communities under artificial selection (AS; on the top) and no artificial selection (NS, on the bottom). The data from 16S RNA gene sequencing provide information at the genus level whereas the data from *gyrB* gene sequencing allow to distinguish the different strains, except for the two *E. coli* strains and the two *P.* sp. ADP strains. The relative frequencies of the different strains in a community were obtained from the sequence counts (note that the 16S RNA gene is a multicopy gene whereas *gyrB* is a monocopy gene). The samples in which a contamination was detected are included in the figure for consistency between the selection treatments.

**Table S1 Deviance table of the analysis of variance (ANOVA) of the optical density (OD) of the ancestors and evolved monocultures and communities under artificial and no artificial selection depending on the initial richness level.** The effect of the initial richness level (1, 2, 4, 8,16), the history (ancestors, evolved under artificial selection, evolved under no artificial selection), the dataset (experimental evolution, post-selection) and their interactions on OD were estimated with a linear mixed model including the identity of the selection unit as a random effect factor on the intercept. The conditional $R^2$ is presented (i.e. variance explained by both fixed and random effect factors; the marginal $R^2$ – fixed effect factors only – was 0.48).

| | Df | Chi squared | p |
|---|---|---|---|
| Initial richness level | 4 | 54.5 | **$4.07 \times 10^{-11}$** |
| History | 2 | 82.6 | **$<2.2 \times 10^{-16}$** |
| Dataset | 1 | 1.18 | 0.278 |
| Initial richness level * History | 8 | 3.01 | 0.934 |
| Initial richness level* Dataset | 4 | 1.45 | 0.836 |
| History * Dataset | 2 | 3.16 | 0.206 |
| Initial richness level * History * Dataset | 8 | 6.08 | 0.639 |
| | | *$R^2$=0.75* | |

**Table S2 Deviance tables of the analyses of variance (ANOVA) of the growth parameters of the ancestors and evolved monocultures and communities under artificial and no artificial selection depending on the initial richness level.** The effect of the initial richness level (1, 2, 4, 8, 16), the history (ancestors, evolved under artificial selection, evolved under no artificial selection) and their interaction on the slope of the exponential growth phase and the time to reach the maximum OD were estimated with a linear mixed model including the identity of the selection unit as a random effect factor on the intercept. The conditional $R^2$ are presented (i.e. variance explained by both fixed and random effect factors; the marginal $R^2$ – fixed effect factors only – were 0.24 and 0.40).

| | Slope | | | Time to maximum OD | | |
|---|---|---|---|---|---|---|
| | Df | Chi squared | p | Df | Chi squared | p |
| Initial richness level | 4 | 12.0 | **1.76x10$^{-2}$** | 4 | 9.14 | 5.77x10$^{-2}$ |
| History | 2 | 16.3 | **2.88x10$^{-4}$** | 2 | 86.6 | **<2x10$^{-16}$** |
| Initial richness level * History | 8 | 5.01 | 0.756 | 8 | 13.2 | 0.107 |
| | | *$R^2$ = 0.56* | | | *$R^2$=0.69* | |

**Table S3 Deviance table of the analysis of variance (ANOVA) of the optical density (OD) of the ancestors and evolved monocultures and communities under artificial and no artificial selection depending on the initial richness level and the substrate category.** The effect of the initial richness level (1, 2, 4, 8, 16), the history (ancestors, evolved under artificial selection, evolved under no artificial selection), the substrate category amines, amino acids, carbohydrates, carboxylic acids, phenolic compounds, polymers) and their interactions on OD were estimated with a linear mixed model including the identity of the selection unit and the substrate as random effect factors on the intercept. The conditional $R^2$ is presented (i.e. variance explained by both fixed and random effect factors; the marginal $R^2$ – fixed effect factors only – was 0.20).

| | Df | Chi squared | p |
|---|---|---|---|
| Initial richness level | 4 | 42.7 | **$1.21 \times 10^{-8}$** |
| History | 2 | 14.7 | **$6.42 \times 10^{-4}$** |
| Substrate category | 5 | 4.17 | 0.526 |
| Initial richness level * History | 8 | 38.5 | **$6.10 \times 10^{-6}$** |
| Initial richness level * Substrate category | 20 | 127 | **$<2.2 \times 10^{-16}$** |
| History * Substrate category | 10 | 12.5 | 0.256 |
| Initial richness level * History * Substrate category | 40 | 29.2 | 0.896 |
| | | *$R^2$=0.58* | |

**Table S4 Theoretical community composition at the beginning of the experiment.** Six different communities per richness level (four levels) were built. The communities from level 16 were built at random from a pool of 18 strains and the communities from the lower levels of richness were subsets of the communities from the higher levels. The full names of the strains are given in Table 1.

| | **C1** | **C2** | **C3** | **C4** | **C5** | **C6** |
|---|---|---|---|---|---|---|
| **Level 16** | *V.* sp. 38R<br>*P. saltans* DSM 12145<br>*A. aminovorans* SR38<br>*P. knackmussii* DSM 6978<br>*M.* sp. C448<br>*C. necator* JMP134<br>*P.* sp. ADP3<br>*E. coli* K12<br>*E. coli* WA803<br>*P.* sp. ADPe<br>*A. eutrophus* JMP131<br>*S. wittichii* RW1<br>*R.* sp.<br>*A.* sp.<br>*A.* sp. BS2<br>*P. azelaica* | *A. aminovorans* SR38<br>*P. knackmussii* DSM 6978<br>*P. saltans* DSM 12145<br>*A. eutrophus* JMP131<br>*P.* sp. ADPe<br>*E. coli* WA803<br>*S. wittichii* RW1<br>*A.* sp. 9023<br>*E. coli* K12<br>*C. necator* JMP134<br>*A.* sp. BS2<br>*M.* sp. C448<br>*D. fermentans* DSM 18053<br>*P.* sp. ADP3<br>*A.* sp.<br>*V.* sp. 38R | *E. coli* WA803<br>*A.* sp. 9023<br>*A.* sp. BS2<br>*P. saltans* DSM 12145<br>*M.* sp. C448<br>*C. necator* JMP134<br>*A. eutrophus* JMP131<br>*S. wittichii* RW1<br>*P. azelaica*<br>*P.* sp. ADPe<br>*A.* sp.<br>*D. fermentans* DSM 18053<br>*E. coli* K12<br>*R.* sp.<br>*P. knackmussii* DSM 6978<br>*A. aminovorans* SR38 | *D. fermentans* DSM 18053<br>*E. coli* K12<br>*P. saltans* DSM 12145<br>*V.* sp. 38R<br>*A. eutrophus* JMP131<br>*M.* sp. C448<br>*P.* sp. ADPe<br>*S. wittichii* RW1<br>*P. knackmussii* DSM 6978<br>*P.* sp. ADP3<br>*R.* sp.<br>*A.* sp. 9023<br>*E. coli* WA803<br>*P. azelaica*<br>*A.* sp.<br>*A.* sp. BS2 | *P.* sp. ADPe<br>*A.* sp. 9023<br>*A.* sp.<br>*S. wittichii* RW1<br>*V.* sp. 38R<br>*P.* sp. ADP3<br>*P. knackmussii* DSM 6978<br>*C. necator* JMP134<br>*E. coli* WA803<br>*P. azelaica*<br>*A. aminovorans* SR38<br>*R.* sp.<br>*P. saltans* DSM 12145<br>*M.* sp. C448<br>*A. eutrophus* JMP131<br>*E. coli* K12 | *A. aminovorans* SR38<br>*A.* sp. 9023<br>*A.* sp. BS2<br>*P.* sp. ADPe<br>*M.* sp. C448<br>*V.* sp. 38R<br>*D. fermentans* DSM 18053<br>*P. azelaica*<br>*C. necator* JMP134<br>*P.* sp. ADP3<br>*P. knackmussii* DSM 6978<br>*R.* sp.<br>*A. eutrophus* JMP131<br>*S. wittichii* RW1<br>*E. coli* K12<br>*P. saltans* DSM 12145 |
| | **C7** | **C8** | **C9** | **C10** | **C11** | **C12** |
| **Level 8** | *V.* sp. 38R<br>*P. saltans* DSM 12145<br>*A. aminovorans* SR38<br>*P. knackmussii* DSM 6978<br>*M.* sp. C448<br>*C. necator* JMP134<br>*P.* sp. ADP3<br>*E. coli* K12 | *E. coli* WA803<br>*P.* sp. ADPe<br>*A. eutrophus* JMP131<br>*S. wittichii* RW1<br>*R.* sp.<br>*A.* sp.<br>*A.* sp. BS2<br>*P. azelaica* | *A. aminovorans* SR38<br>*P. knackmussii* DSM 6978<br>*P. saltans* DSM 12145<br>*A. eutrophus* JMP131<br>*P.* sp. ADPe<br>*E. coli* WA803<br>*S. wittichii* RW1<br>*A.* sp. 9023 | *E. coli* K12<br>*C. necator* JMP134<br>*A.* sp. BS2<br>*M.* sp. C448<br>*D. fermentans* DSM 18053<br>*P.* sp. ADP3<br>*A.* sp.<br>*V.* sp. 38R | *E. coli* WA803<br>*A.* sp. 9023<br>*A.* sp. BS2<br>*P. saltans* DSM 12145<br>*M.* sp. C448<br>*C. necator* JMP134<br>*A. eutrophus* JMP131<br>*S. wittichii* RW1 | *P. azelaica*<br>*P.* sp. ADPe<br>*A.* sp.<br>*D. fermentans DSM 18053*<br>*E. coli* K12<br>*R.* sp.<br>*P. knackmussii* DSM 6978<br>*A. aminovorans* SR38 |
| | **C13** | **C14** | **C15** | **C16** | **C17** | **C18** |
| **Level 4** | *V.* sp. 38R<br>*P. saltans* DSM 12145<br>*A. aminovorans* SR38<br>*P. knackmussii* DSM 6978 | *M.* sp. C448<br>*C. necator* JMP134<br>*P.* sp. ADP3<br>*E. coli* K12 | *E. coli* WA803<br>*P.* sp. ADPe<br>*A. eutrophus* JMP131<br>*S. wittichii* RW1 | *R.* sp.<br>*A.* sp.<br>*A.* sp. BS2<br>*P. azelaica* | *A. aminovorans* SR38<br>*P. knackmussii* DSM 6978<br>*P. saltans* DSM 12145<br>*A. eutrophus* JMP131 | *P.* sp. ADPe<br>*E. coli* WA803<br>*S. wittichii* RW1<br>*A.* sp. 9023 |
| | **C19** | **C20** | **C21** | **C22** | **C23** | **C24** |
| **Level 2** | *V.* sp. 38R<br>*P. saltans* DSM12145 | *A. aminovorans* SR38<br>*P. knackmussii* DSM 6978 | *M.* sp. C448<br>*C. necator* JMP134 | *P.* sp. ADP3<br>*E. coli* K12 | *E. coli* WA803<br>*P.* sp. ADPe | *A. eutrophus* JMP131<br>*S. wittichii* RW1 |
